# Effective connectivity predicts distributed neural coding of perceptual decision confidence, uncertainty and speed

**DOI:** 10.1101/2024.03.09.584217

**Authors:** Abdoreza Asadpour, KongFatt Wong-Lin

**Author notes:** Abdoreza Asadpour is now with the School of Life Sciences, University of Sussex, UK. Corresponding authors: Abdoreza Asadpour; KongFatt Wong-Lin.

## Abstract

Decision-making is often accompanied by a level of confidence regarding the accuracy of one’s decision. Previous studies have indicated neural activity associated with perceptual decision confidence during sensory stimulus presentation. Choice-based reaction time (RT) has been suggested as an indirect but more objective measure of decision confidence – generally faster RT for higher confidence. However, it is unclear whether choice confidence and RT are mediated by distinct neural pathways, and whether their neural correlates are encoded nonlinearly. Within a perceptual decision-making task, we applied fMRI-informed EEG-based effective connectivity analysis via dynamic causal modelling (DCM) on event-related potentials and found the frontoparietal network for fast-vs-slow RT condition to be different from that of high-vs-low confidence rating condition. Furthermore, trial-by-trial DCM analysis predicted cortical layer-based, distributed and nonlinear coding of RT, confidence or uncertainty. Collectively, our study suggests that decision confidence and speed are instantiated by different dynamical networks distributed across cortical layers.

## Introduction

Perceptual decision confidence is the internal evaluation of the accuracy of one’s perceptual decision (Fleming, 2021; Metcalfe & Arthur P. Shimamura, 1994; Pleskac & Busemeyer, 2010). Studies have identified the neural correlates of perceptual decision confidence during the early phase of stimulus presentation (Gherman & Philiastides, 2015, 2018; Grogan et al., 2023; Hebart et al., 2016; Heereman et al., 2015; Hoven et al., 2022; Li & Yang, 2012; Zizlsperger et al., 2014). In particular, neuroimaging studies have revealed the involvement of several brain regions in perceptual decision-making and confidence evaluations, including the frontal, parietal, temporal, and occipital lobes, as well as the insula and cerebellum (Gherman & Philiastides, 2015, 2018; Hebart et al., 2016; Heereman et al., 2015; Hoven et al., 2022; Li & Yang, 2012; Zizlsperger et al., 2014) (Table S1). Computational models have suggested the monitoring of perceptual decision confidence or uncertainty to affect or control decision speed via common neural pathways (Atiya et al., 2019; Desender et al., 2019) (Fig. 1A, H1). However, there may exist distinct neural pathways for decision confidence and choice-based response times (RTs), as illustrated in Fig. 1A, H2.

**Fig. 1.**
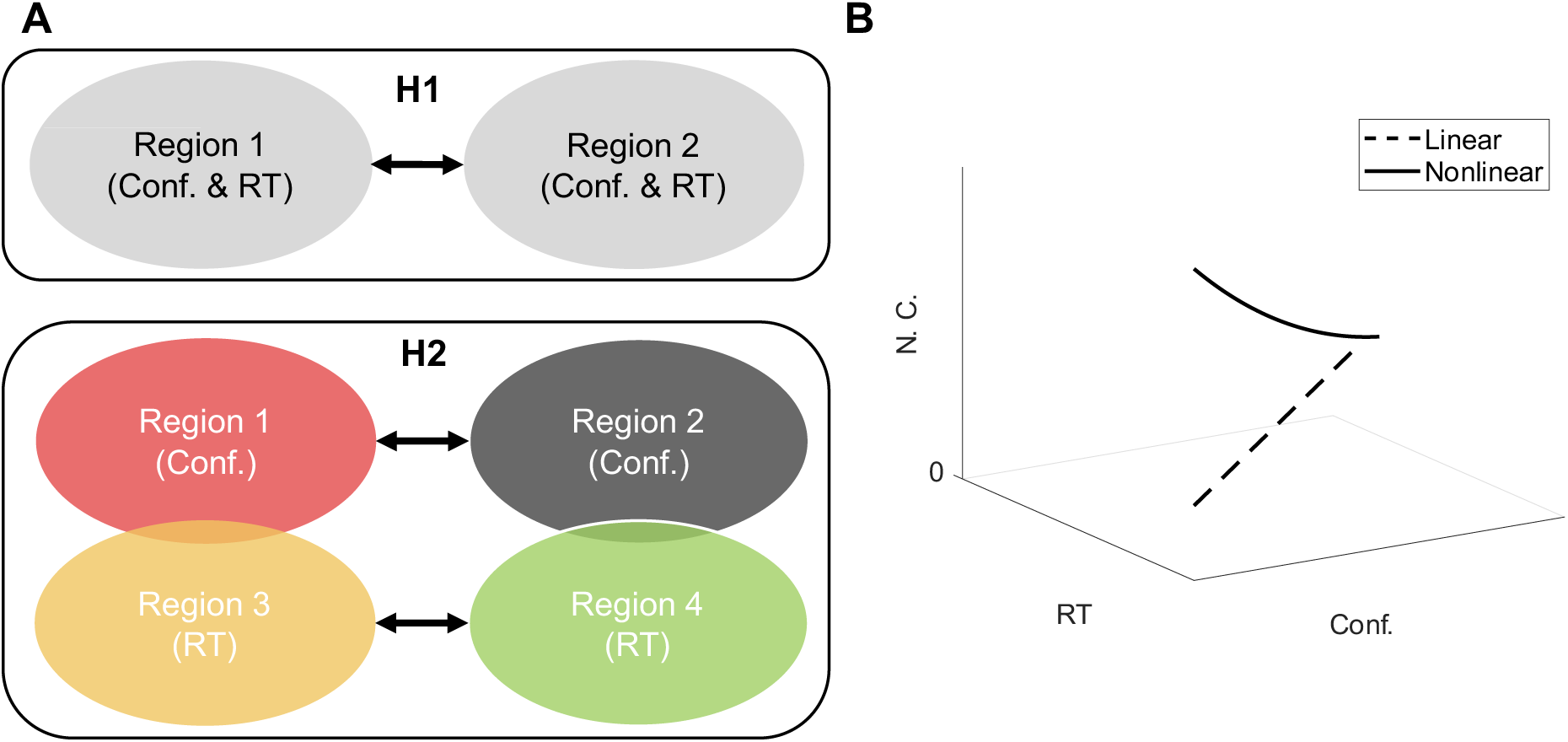
Hypotheses for neural encoding of decision confidence and speed. (**A**) Two competing hypotheses on the neural pathways for subjective confidence (Conf.) and reaction time (RT) during a perceptual confidence task; H1: Subjective confidence and choice-based RTs share the same neural pathway H2: Subjective confidence and RT have distinct pathways with overlaps. (**B**) Possible relationships of neural correlates (N. C.) with confidence and choice-based RT during a perceptual decision confidence task.

While functional magnetic resonance imaging (fMRI) and electroencephalography (EEG)-informed fMRI have been the primary tools for identifying active brain regions and examining functional connectivity (Murta et al., 2015), directed connectivity approaches such as dynamic causal modelling (DCM) can offer invaluable insights into the causal relationships between these regions through explicit neural circuit modelling (Stephan et al., 2007). Existing studies employing DCM have elucidated the effective connectivity between brain regions during various perceptual decision-making tasks, revealing specific directed connections between frontal, parietal, and temporal regions that are critical for decision-making processes (Brázdil et al., 2007; Dowlati et al., 2016; Lamichhane & Dhamala, 2015; Nguyen et al., 2014; Raffin et al., 2022; Tsumura et al., 2022). However, there is limited investigation using DCM for understanding decision confidence, with the only study focusing on the neural dynamics underlying belief updating and confidence in the dorsolateral prefrontal and anterior cingulate cortices (Fiore & Gu, 2022). Hence, there is a notable gap in our understanding of the directed information flow and neural circuit dynamics of subjective perceptual decision confidence.

To complement the investigation of subjective decision confidence evaluation, which can vary widely across participants (Desender et al., 2018), previous studies have sought more objective measures of decision confidence, particularly using choice-based RT such that faster RTs are generally associated with higher confidence levels (Pleskac & Busemeyer, 2010; Vickers, 1979). Prior research employing EEG and fMRI has illuminated the neural correlates of confidence and decision-making processes, demonstrating activations in regions such as the prefrontal, parietal, temporal and occipital cortices in relation to subjective confidence judgements and RTs (Boldt et al., 2019; Fleming & Dolan, 2012; Gherman & Philiastides, 2018; Hebart et al., 2016; Philiastides & Ratcliff, 2013; Sadras et al., 2023). These findings, including fMRI and EEG source analyses, demonstrate a relationship between brain regions, with early encoding of confidence and decision variables. They reveal a complex network that is synthesising sensory evidence and performing evaluations to generate both decisions and levels of confidence. In addition, these studies mostly assume a linear relationship of neural correlates with RT and confidence specifically in fMRI data analysis and therefore they performed a GLM approach to unravel this relationship (Fig. 1B, dashed line). However, there is evidence suggesting that the relationship between neural correlates and confidence may be monotonic and nonlinear, a possibility that remains underexplored but has been reported by <u>Boldt & Yeung (2015)</u>. Moreover, the causal relationships between brain regions involved in confidence judgements and the temporal dynamics of these processes remain underexplored, particularly in investigating the potential nonlinear interactions between neural correlates and behaviour (Fig. 1B, solid line). The roles across cortical areas are also unclear. These can be delineated with an integration of these modalities using DCM approach with source-EEG dynamics (David et al., 2006) while peering inside the neural model.

In this work, we investigate the complex relationship between neural activities, subjective confidence ratings, and choice-based RTs to delineate their collective impact on confidence evaluation processes, and whether perceptual decision confidence and speed share similar neural circuits. We employed DCM to analyse an open EEG-fMRI dataset (Gherman & Philiastides, 2020), using an fMRI-informed EEG approach (Penny et al., 2011). We also applied an innovative trial-by-trial DCM approach, informed by the averaged DCM, to predict nonlinear relationships between estimated neural population activities, decision confidence and choice-based RT. Our work identified not only the neural correlates of decision confidence, consistent with previous studies (Gherman & Philiastides, 2015, 2018; Hebart et al., 2016; Heereman et al., 2015; Hoven et al., 2022; Li & Yang, 2012; Zizlsperger et al., 2014), but also for choice-based RT. Crucially, these two measures were associated with distinct brain networks. Furthermore, the DCM model predicted distributed and nonlinear neural coding of choice confidence, uncertainty and RT, with some within deeper cortical regions, a finding that cannot be captured by scalp-level EEG analysis, while challenging the prevailing assumption of shared neural pathways for decision confidence and speed.

## Methods

We utilised an open concurrent EEG-fMRI dataset on perceptual decision confidence to investigate the active brain regions and effective connectivity of fMRI-informed EEG data (Gherman & Philiastides, 2020). The dataset comprised 24 participants aged 20–32 years; who discriminated the direction of coherent motion in random-dot kinematogram (RDM) (Britten et al., 1992) and rated their confidence. However, due to inconsistencies in the structural and functional data of the last participant, analyses were conducted on the remaining 23 participants.

### Experimental Paradigm

The perceptual decision task involved visually discriminating the direction of coherent motion in an RDM. The RDM stimuli consisted of white dots moving on a black background within a circular aperture, with a subset moving coherently to form the signal while the rest moved randomly. The difficulty was controlled by adjusting the proportion of coherently moving dots, aiming to maintain an overall performance of approximately 75% correct responses. Each trial began with an RDM stimulus presented for up to 1.2 seconds, during which participants made a left or right-directional discrimination using a button press. This was followed by a blank screen and a random delay of 1.5 to 4 seconds. Subsequently, participants rated their confidence on a horizontal bar scale for 3 seconds. Trials ended with another random delay of 1.5 to 4 seconds. Each participant performed two experimental blocks of 160 trials each, corresponding to two separate fMRI runs. All behavioural responses were executed using the right hand on an MR-compatible button box (Gherman & Philiastides, 2018). Fig. 2A shows the sequence within a trial.

**Fig. 2.**
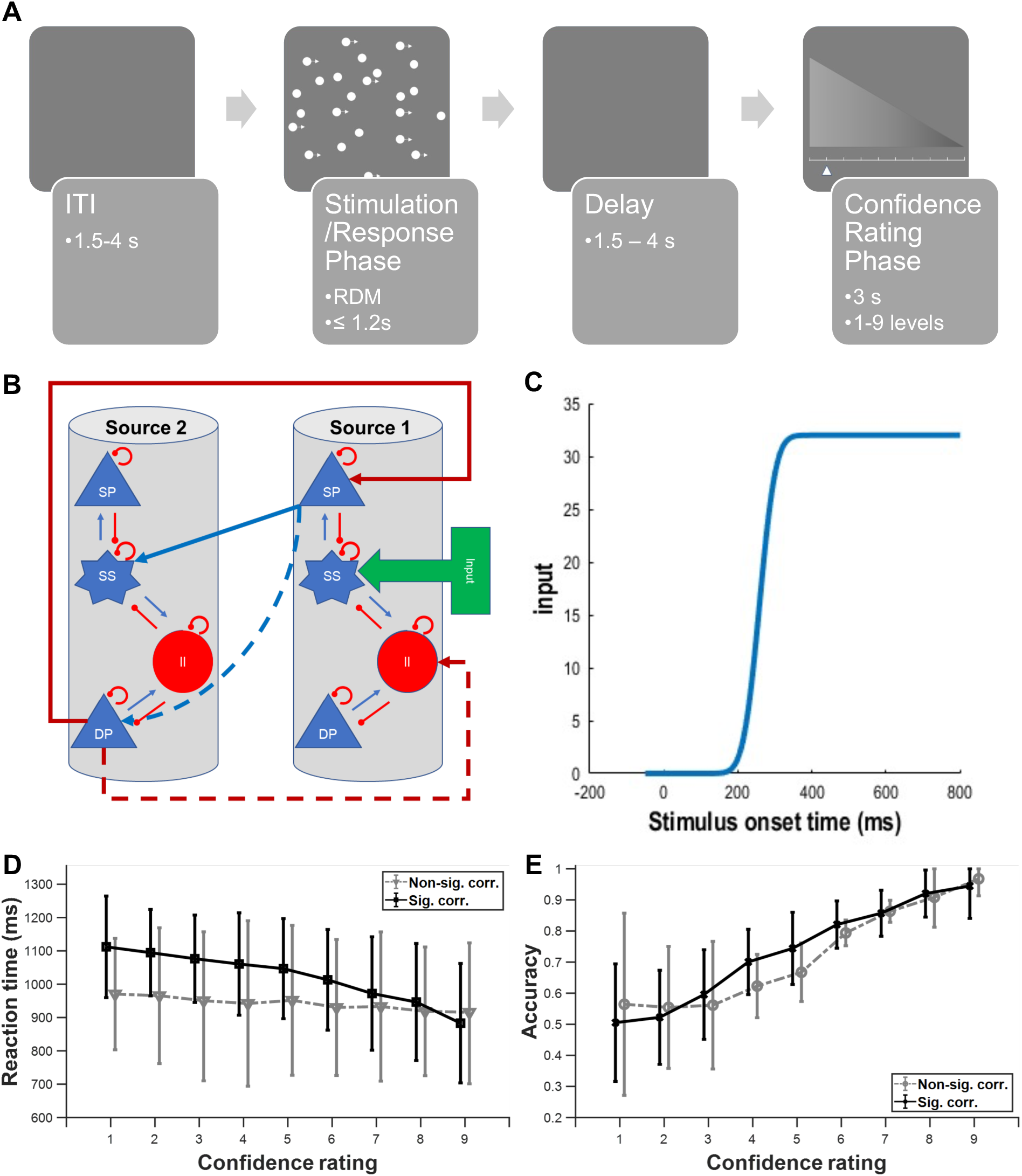
Effective connectivity analysis of a motion discrimination task with confidence rating and RT. (**A**) Schematics of data acquisition protocol (Gherman & Philiastides, 2018). Task begins with a variable intertrial interval (ITI) followed by presentation of a random-dot kinematogram and participants were instructed to respond based on their perceived motion coherence direction (stimulation/response phase). After a variable delay period, the confidence rating phase appears for 3 s in which participants have to rate their confidence level (1 to 9) of their decision just made. (**B**) Example model in DCM. Internal and external connections of two cortical columns based on the canonical microcircuit model (CMC); four distinct neural populations per cortical column: excitatory spiny stellate (SS) neural population, inhibitory interneurons (II), deep excitatory pyramidal (DP) neural population, and superficial excitatory pyramidal (SP) neural population. Internal blue (red) connections: excitatory (inhibitory) connections. External blue (red) arrows between sources: forward (backward) connections. (**C**) Cumulative Gaussian input with onset at 200 ms after actual stimulation. (**D**) Distribution of RTs for different self-reported confidence levels across all participants, excluding the last participant. Black: significant Spearman’s correlation (p<0.05) between confidence level and RTs; grey: non-significant correlations (**E**) Choice accuracy of trials for different reported confidence levels across all participants. Black: significant Spearman’s correlation (p<0.05) between confidence level and RTs; grey: non-significant correlations.

### Data Acquisition

Functional data were acquired using a T2*-weighted gradient echo echo-planar imaging (EPI) sequence on a 3-T Siemens MRI scanner. The parameters included 32 interleaved slices, a 0.3 mm gap, 3 × 3 × 3 mm voxel size, 70 × 70 matrix size, 210 mm field of view (FOV), 30 ms echo time (TE), 2000 ms repetition time (TR), and 80° flip angle. A high spatial resolution anatomical volume was obtained using a T1-weighted sequence (Gherman & Philiastides, 2018).

Simultaneously, EEG data were collected using a 64-channel MR-compatible system from Brain Products, Germany, with a 5000 Hz sampling rate. Electrodes were positioned according to the 10 – 20 system, with additional nasion reference and ground electrodes. Data acquisition was synchronised with the MRI scanner (Gherman & Philiastides, 2018).

### Data preprocessing

The fMRI data preprocessing included slice-timing correction, high-pass filtering with a cutoff of 100 seconds, spatial smoothing using an 8 mm Gaussian kernel, and head motion correction. SPM12 (Penny et al., 2011) was used for realignment, extraction of motion parameters, co-registration of the mean fMRI volumes with the structural data, and normalisation to standard brain space.

EEG data preprocessing was conducted using MATLAB. Gradient artefacts were corrected by subtracting average artefact templates, followed by a 12 ms median filter. A 0.5 – 40 Hz band-pass filter was applied, and data were downsampled to 1000 Hz. Eye movement and cardiac-related artefacts were minimised using principal component analysis, with baseline correction performed by removing the average signal during the 100 ms prestimulus interval (Gherman & Philiastides, 2018). No additional preprocessing steps were applied in our study.

### fMRI Data Analysis

We employed the generalised linear model (GLM) analysis to statistically analyse the functional data and extract active brain regions under different conditions (high vs. low confidence ratings and fast vs. slow RTs). SPM generates the fMRI time series by convolving a time series of delta functions representing the event onsets with a hemodynamic response function (HRF) (Penny et al., 2011). Standard HRF utilised extensively in SPM is a mixture of two or more gamma functions (Penny et al., 2011).

RTs were clustered into four groups using k-means clustering (MacQueen, 1967), performed separately for each participant. RTs below the second cluster centre were classified as fast, those between the second and third cluster centres as medium, and those above the third cluster centre as slow. Although reaction times (RT) were categorized into three levels—fast, medium, and slow—our analysis focused on fast vs. slow RTs to provide a clearer contrast between the extremes. Similarly, for confidence levels, we concentrated on high vs. low confidence ratings for the same reason, with intermediate levels excluded from the primary analysis.

Then, first-level analysis was conducted within each participant using T-contrast calculations to compare conditions (high vs. low confidence ratings and fast vs. slow RTs). This involved calculating the t-statistic for each brain voxel to discern differences between the means of two conditions. P-values were adjusted using Bonferroni correction and family-wise error (FWE) correction at a level of 0.05. Second-level analysis pooled data from all participants to facilitate group-level T-contrast analysis, comparing epochs of high vs. low confidence ratings and fast vs. slow RTs.

### DCM Analysis

DCM was used to explore the directed connectivity between brain regions. Both averaged event-related potential (ERP)-DCM and trial-by-trial ERP-DCM approaches were employed. We created the model space based on the T-contrast analysis of fMRI data separately for high/low confidence and fast/slow reaction time (RT) conditions to ensure parsimonious models. Combining regions from different conditions or extracting active regions against baseline to create the model space would result in more complex models, which the DCM approach tends to favour due to their increased complexity (Litvak et al., 2019). The canonical microcircuit (CMC) neural mass model was selected, which includes excitatory spiny stellate (SS) neural population, inhibitory interneurons (II), deep excitatory pyramidal (DP) neural population, and superficial excitatory pyramidal (SP) neural population (Bastos et al., 2012; Pinotsis et al., 2013). A simplified illustration of these relationships is provided here (Fig. 2B)

#### Averaged ERP-DCM

For each participant and condition, we averaged the epochs corresponding to high and low confidence ratings as well as fast and slow RTs from the stimulation phase, which spanned from 50 ms prior to stimulus onset to 1.2 seconds after onset. However, our DCMs estimated neural activity until 800 ms post-stimulus onset to particularly investigate early neural correlates.

We selected the CMC model due to its suitability for modelling higher brain frequencies, including gamma activity, which are relevant to perceptual decision-making processes (Donner et al., 2009). The cumulative Gaussian signal was used as the DCM stimulus input, with onset at 200 ms post-stimulus to account for the delay due to sensory transduction (FitzGerald et al., 2015) (Fig. 2C). The estimated neural activity was linearly mapped to scalp EEG activities through a cortical surface patch (Penny et al., 2011).

#### Trial-by-Trial ERP-DCM

Following the averaged ERP-DCM analysis, we employed a trial-by-trial ERP-DCM approach to analyse and compare the neural dynamics under different conditions (high vs. low confidence ratings and fast vs. slow RTs) during the stimulation phase. This approach aimed to uncover trial-by-trial variability and specific neural interactions related to early confidence evaluation.

For each trial, we used the data from the entire epoch, ranging from -50 ms to 1.2 seconds from the stimulation onset, to train the DCMs. The initial parameters of the trial-by-trial DCMs were kept the same as those used for the winning models obtained in the averaged ERP-DCM analysis for each condition.

### Statistical Analysis of averaged ERP-DCM

#### Bayesian Model Selection

Bayesian model selection (BMS) was employed to identify the most likely model from the set of candidate models. BMS compares different models based on their likelihood given the observed data. The random effects (RFX) procedure was used, which accounts for variability between participants, assuming different performance mechanisms for different participants (Penny et al., 2011).

BMS calculates the model evidence, which is the probability of the data given the model, integrating over all possible parameter values. The model with the highest evidence is considered the most probable. We also employed Bayesian model averaging (BMA) to account for model uncertainty by averaging over models weighted by their posterior probabilities (Trujillo-Barreto et al., 2004).

#### Parametric Empirical Bayes

To investigate variability between participants, we employed the Parametric Empirical Bayes (PEB) method on participant-specific effective connectivity parameters retrieved from the winning DCM. PEB is a hierarchical Bayesian approach that provides participant-level estimated connection strengths and their uncertainty at the group level (Penny et al., 2011).

### Statistical Analysis of trial-by-trial ERP-DCM

Support vector regression (SVR) with linear and nonlinear (Gaussian) kernels was utilised to establish associations between neural activity patterns and behavioural metrics (Drucker et al., 1996). In particular, we were interested in investigating the (non)linear relationship and the importance of each time point during the stimulation phase in determining behavioural metrics (Fig. 1B). SVR models were trained for each participant for each condition using the estimated neural population activities for each trial corresponding to each condition.

Permutation importance was used to assess the significance of each feature in predicting behavioural outcomes (Breiman, 2001). Additionally, we conducted sensitivity analysis by perturbing the data to determine if the relationship between neural population activity and behavioural data (RT or confidence rating) is positive or negative (Hinch, 1991). Next, bootstrapping was performed to construct distributions of predictive performances, determining significance by calculating confidence intervals through resampling (Efron, 1979).

For comprehensive methodological details, please refer to Supplementary Material 1.

## Results

### A perceptual decision task with negative correlation between decision confidence rating and RT

In our analysis and modelling, we utilised an open concurrent EEG-fMRI dataset on human perceptual decision confidence. The task involved visually discriminating the direction of coherent motion in RDM (Britten et al., 1992) in which a subset of dots moved coherently in certain direction while the rest moved randomly. The difficulty of the task was manipulated by adjusting the proportion of coherently moving dots, aiming to maintain an overall performance of approximately 75% correct responses. The protocol consisted of a stimulation/response phase in the presence of the RDM (extinguished upon motor reporting of choice by clicking a mouse button), followed by a variable delay period, before a subjective confidence rating phase, and then a variable intertrial interval (ITI) (Fig. 2A). For a detailed description of the protocol and task, please refer to the Methods section, Supplementary Material 1, and the original study (Gherman & Philiastides, 2018).

Our study’s primary aim is to uncover the early neural correlates of subjective confidence evaluations and choice-based RTs (serving as a more objective confidence measure) in this perceptual decision-making task. To this end, we employed Spearman’s correlation to investigate the relationship between subjective confidence ratings and RTs, with faster RTs generally being associated with higher confidence levels (Pleskac & Busemeyer, 2010; Vickers, 1979). Subjects who did not show a significant monotonic correlation between reaction time (RT) and confidence were excluded from the primary DCM analysis. This exclusion was made to ensure that the data used for modelling reflected participants who engaged with the task consistently. Our findings revealed that 17 of the 23 participants exhibited a significant correlation between their RTs and subjective confidence ratings (p<0.05) (Fig. 2D and E). Additionally, Spearman’s correlation between the delay period and the confidence ratings at the trial level was 0.09 (p<0.001) showing that the delay period did not systematically influence the confidence ratings reported by the participants. We also observed significant differences (p<0.01) in mean accuracy between fast (0.87 ± 0.09) and slow RT (0.69 ± 0.14) trials using the Wilcoxon signed-rank test (Wilcoxon, 1945), which may reflect variations in task engagement or decision strategy. Therefore, we used the data of those 17 participants for further DCM analysis and modelling.

We then proceeded to explore the neural correlates of perceptual decision confidence evaluation through a combination of ERP and fMRI-informed effective connectivity in the form of DCM (David et al., 2006) (see example neural model in Fig. 2B). For fMRI, averaged DCM, and trial-by-trial DCM analyses, we compared the data across two scenarios: contrasting high versus low subjective confidence ratings and comparing fast versus slow RTs as a measure of objective confidence during the stimulation/response phase.

### Averaged ERP-based dynamic causal modelling

To investigate the overall directed connectivity of brain regions involved in confidence evaluation, we employed an averaged ERP-DCM approach, beginning with the analysis of the blood oxygenation level dependent (BOLD)-fMRI data to extract active brain regions under the abovementioned conditions (Methods and Supplementary Material 1). Then, we performed the averaged ERP-DCM analysis to decipher the neural dynamics and network mechanisms underlying perceptual confidence and decision-making processes. To model both gamma-band activities and lower frequency-band activities, given their relevance to perceptual decision-making processes (Donner et al., 2009), we selected the CMC neural mass model (Pinotsis et al., 2013), consisting of four distinct neural populations per cortical column, namely, SS neural population, II, DP neural population, and SP neural population (Bastos et al., 2012; Pinotsis et al., 2013) (Methods and Supplementary Material 1). Fig. 2B illustrated an example with two canonical cortical columns with intrinsic and extrinsic forward and backward connections.

We focused on the estimated averaged neural activity from 50 ms prior to stimulus onset until 800 ms post-stimulus onset. We implemented a cumulative Gaussian signal as our DCM stimulus input, similar to the approach used in FitzGerald et al. (2015), with onset at 200 ms post-stimulus (Fig. 2C) to account for the delay due to sensory transduction (Lamme & Roelfsema, 2000).

### Enhanced estimated frontoparietal cortical source activities and feedback connectivity with higher confidence rating

In our analysis, we compared fMRI-informed EEG-DCM between trials with high and low confidence ratings during the stimulation phase. Confidence levels were categorised into high (≥7), medium (5, 6), and low (≤4) based on participants’ ratings. These rating bands were selected to optimise the number of trials per band for statistical power.

By performing a T-contrast analysis of the fMRI data using the GLM (Penny et al., 2011), we identified three regions in the parietal cortex, including the left precuneus (lPreCUN), left inferior parietal lobule (lIPL), and left superior parietal lobule (lSPL), as well as four regions in the frontal cortex, including the middle frontal gyrus (MFG) and superior frontal gyrus (SFG) in both hemispheres (Figs. S1 and S2). Based on these extracted regions, we created a model space consisting of two likely models (Fig. 3A). We conducted an additional fMRI analysis excluding participants with a nonsignificant relationship between RT and subjective confidence. For the high vs. low confidence comparison during the stimulation phase, no active brain regions were identified, highlighting the decreased statistical power of fMRI analysis, specifically by using FWE correction, when certain participants were excluded (Figs. S3 and S4). We then tested the brain activations of all the participants during the stimulation phase against the implicit baseline. This analysis showed that the active areas include undesired regions related to motor movements, sensory stimulus processing, and other non-relevant brain activities (Fig. S5), obscuring our ability to extract key regions specifically relevant to confidence or choice-based RT. We also examined other scenarios for correct versus error trials and right versus left choices but did not find any significant active region (Figs. S6, S7, S8, and S9).

**Fig. 3.**
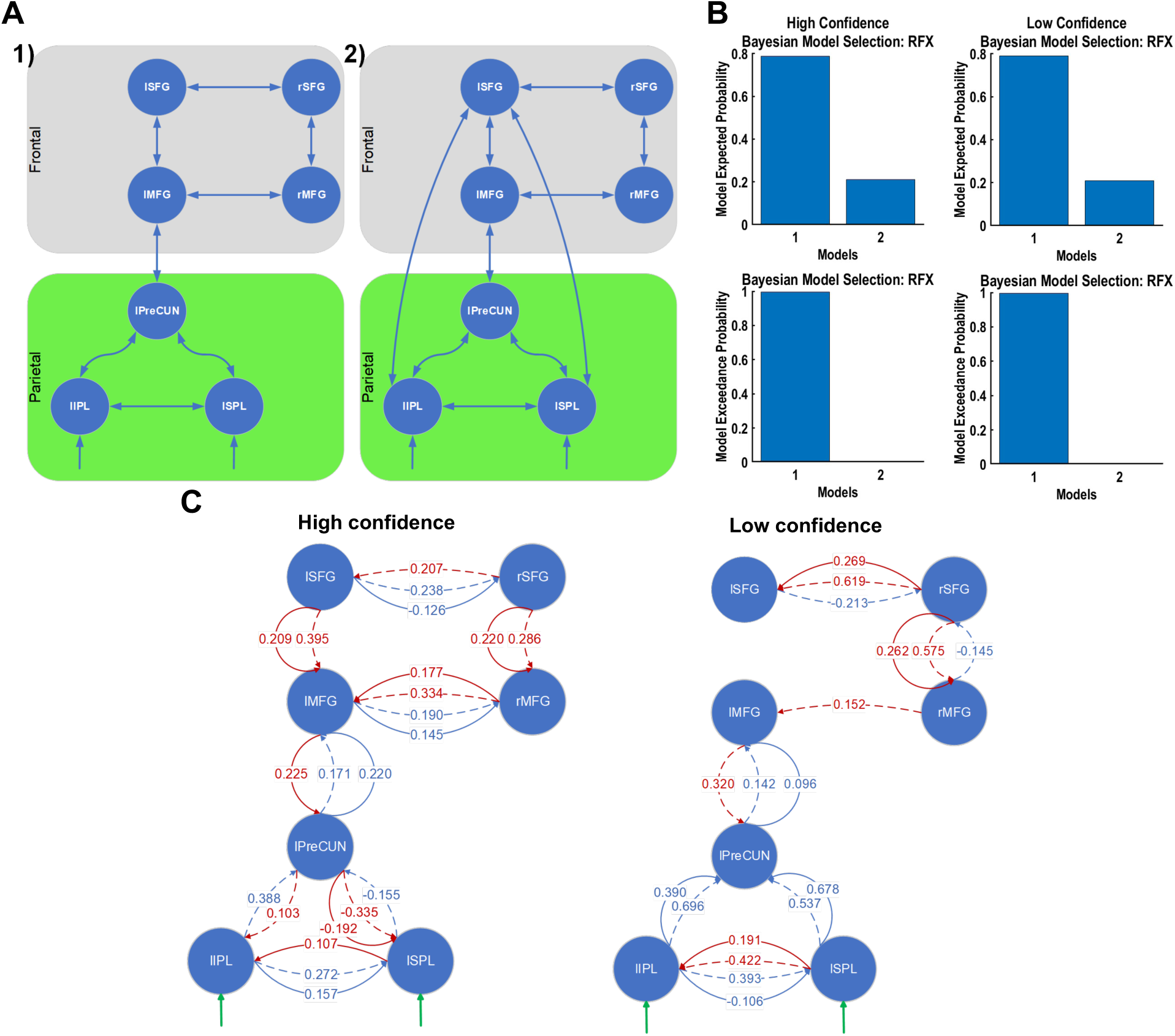
More frontoparietal connections with higher confidence but stronger parietal connectivity with lower confidence. (**A**) Model space for high-vs-low confidence rating conditions in the stimulation phase. Model space for high-vs-low confidence contains plausible connections between active BOLD-fMRI based brain (including parietal (green) and frontal (grey)) regions. (**B**) Bayesian model selection using random-effect approach. Model 1 was the winning model. (**C**) Forward (blue) and backward (red) connections in the winning model (1) with more than 75% probability of occurrence among participants using the PEB approach.

These two models were selected to specifically assess the role of direct frontoparietal connections in the neural mechanisms underlying confidence. The first model comprised plausible connections between regions, reflecting the hypothesized information flow within the frontoparietal network during confidence judgments. The second model extended the first by adding a direct parietal-to-frontal cortex connection from the lIPL and lSPL to lSFG, to test the hypothesis that direct frontoparietal connections contribute significantly to the generation of confidence judgments (Heekeren et al., 2008; Summerfield & Tsetsos, 2015). In addition to our two models, we explored the model space further by conducting a preliminary Bayesian model reduction (BMR) (Penny et al., 2011) on a fully connected model. This analysis revealed that the resultant reduced model, while statistically simplified due to fewer parameters and connections, might lack biological plausibility because of sparse connections between regions, especially the absence of connections between parietal and frontal cortices (Jung et al., 2017). Despite this, we compared the reduced model against our two initially proposed models using BMS (Penny et al., 2011). This comparison confirmed that the models within our current model space were the winning models and suitable for further analysis.

BMS with random effects (Penny et al., 2011) revealed a greater exceedance probability for the first model in both conditions (Fig. 3B). BMA (Penny et al., 2011) in combination with PEB (Penny et al., 2011) showed that the forward connections of SP-to-SS and SP-to-DP (>75% probability of occurrence) for the winning model were stronger between the parietal regions with lower confidence (Fig. 3C). In contrast, these connections had greater connection strength between the lPreCUN and lMFG with higher confidence (Fig. 3C). Other connections showed similar strengths for both conditions. Additionally, for the higher confidence condition, there were more DP-to-SP and DP-to-II backward connections in the winning model, suggesting a greater integration of frontoparietal network during high confidence judgments.

We next used the mean squared error (MSE) to select the best-fitted participant by comparing predicted and observed EEG scalp activity across electrodes. The mean MSE across participants was 0.0039 (0.0052), with a standard deviation of 0.0013 (0.0018) for high (low) confidence condition, indicating a high agreement between the modelled and observed EEG data across the cohort. By visually inspecting the winning model with the best-fitted participant and found that the DCM-based projected EEG activity on the scalp readily recapitulated that of the observed scalp EEG activity over key epochs in a trial for both high and low confidence rating conditions (Fig. S10), albeit some minor differences for the low confidence condition. Moreover, there was a centroparietal positivity (CPP) 370 ms after stimulus onset which converted into a left hemispheric positivity and a right hemispheric negativity at 700 ms. Interestingly, for the low confidence condition, the CPP at 370 ms changed to positivity at 550 ms. This finding is consistent with previous research that has identified the CPP as a neural correlate of the accumulation of evidence during perceptual decision-making processes (Kelly & O’Connell, 2013; O’Connell et al., 2012; Twomey et al., 2015).

For excitatory neural activities, we first analysed those of SP neural populations due to their proximity with scalp EEG (Brown & Friston, 2012). Source space analysis revealed greater activities (averaged scaled voltages) for SP for higher confidence in the parietal regions, including the lIPL, lSPL and lPreCUN, when averaged over all participants (Fig. 4). However, this was not the case for the best-fitted participant (Fig. 4); participant-wise variability. In both high and low confidence conditions, the frontal regions exhibited similar activities, with the exception of the left and right MFG (lMFG and rMFG), where the SP activities were on average more negative with lower confidence (Fig. 4) despite the predominantly positive afferent connections (Fig. 3C). The latter may potentially be encoding decision uncertainty. The lMFG showed differential timescales for low and high confidence; high confidence had an earlier negativity peak (∼180 ms) than low confidence (∼300 ms).

**Fig. 4.**
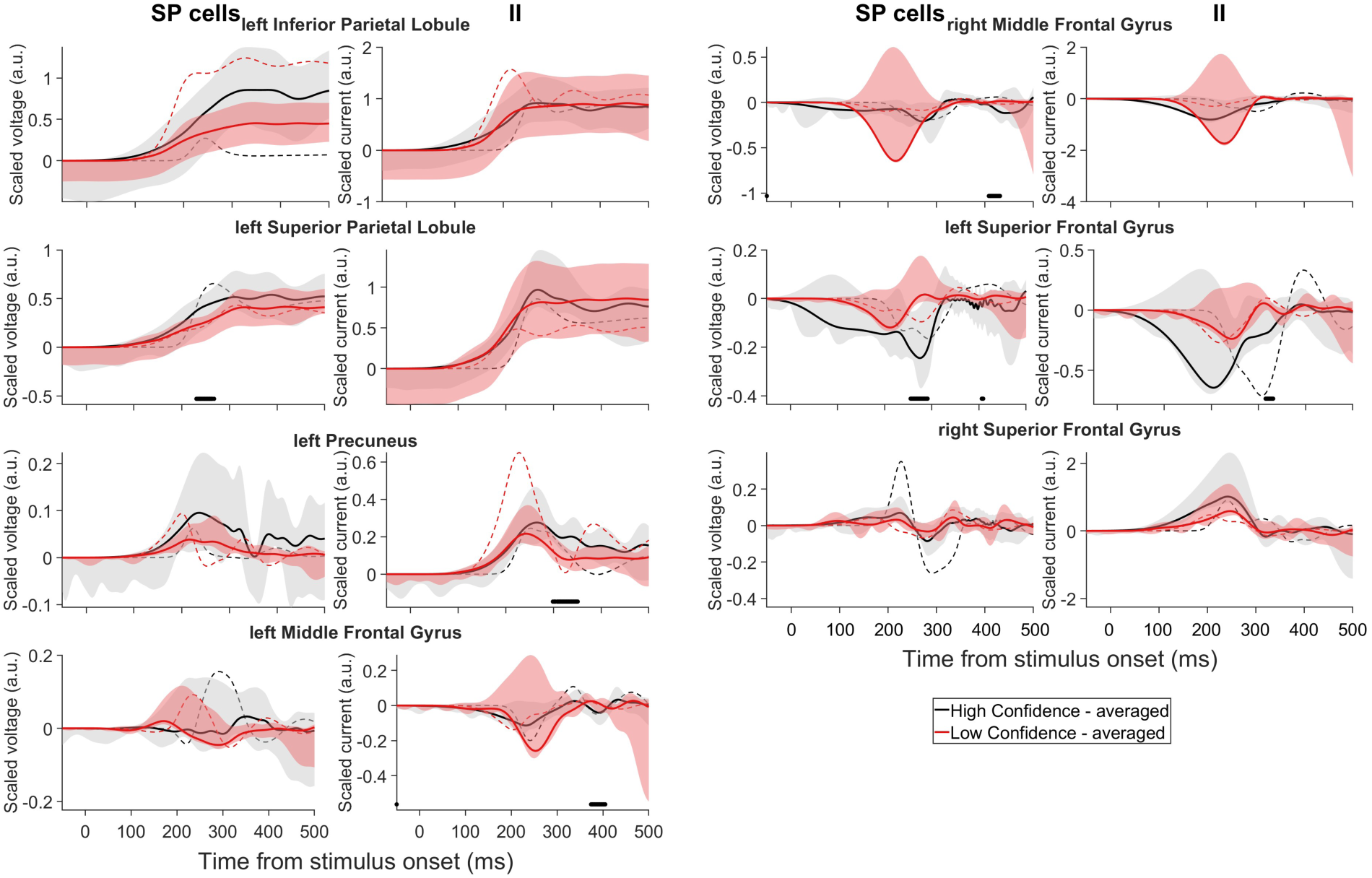
Higher decision confidence generally associated with higher projected source activities. Estimated averaged scaled voltages of SP neural populations and scaled currents of II neural populations of winning model. Black (red): high (low) confidence rating. Solid lines: averaged over participants; dashed lines: best-fitted participant. Shaded regions: 95% confidence intervals. Filled points above x-axis: time points with significant differences between averaged conditions (p<0.05).

The activities (averaged scaled currents) of II neural populations (CMC models in DCM represent scaled II activity as scaled current rather than scaled voltage) displayed similar patterns, apart from the right SFG (rSFG), where the averaged activity was higher with higher confidence. We also found a predominant encoding of subjective confidence by DP neural population in the lPreCUN and potential encoding of subjective uncertainty by SS neural population in the left SFG (lSFG) (Fig. S11).

In summary, by comparing high and low self-reported confidence in perceptual decisions, we found that generally, there were more frontoparietal circuit connections for higher decision confidence, while connections within the parietal cortices were stronger with lower confidence. Additionally, most of the estimated source activities were higher with higher decision confidence, except for the MFG, which may potentially encode decision uncertainty. The inhibitory neural activities were similar to the excitatory ones, except for rSFG, where the inhibitory neural activities were more positive with higher decision confidence. This analysis reveals that the neural mechanisms underlying perceptual decision confidence are associated with distinct patterns of frontoparietal connectivity and neural population activities, providing new insights into the neural basis of confidence in perceptual decision-making. We will next delve into the analysis based on choice-based RT, acting as a more objective measure of decision confidence, exploring how these temporal dynamics align with neural activities.

### Enhanced estimated frontal cortical source activities for faster RTs but stronger parietal cortical connectivity for slower RTs

During the stimulation phase, choice-based RTs were clustered into four groups using k-means clustering (MacQueen, 1967), performed separately for each participant (Methods and Supplementary Material 1). The RTs below the second cluster centre were classified as fast (presumably linked to higher objective decision confidence), and those above the third cluster centre as slow (lower objective decision confidence). Statistical analysis of BOLD-fMRI data using T-contrast between trials with fast and slow RTs revealed five significant active brain regions. Three of these regions were located in the parietal cortex, namely, the right SPL (rSPL), lPreCUN, and right supramarginal gyrus (rSMG), while the remaining two were in the frontal cortex, namely, the left precentral gyrus (lPreCG) and left medial frontal gyrus (lMeFG) (Figs. S12 and S13). Compared to the active brain regions extracted for the high and low subjective decision confidence, where the active regions were the lPreCUN, lIPL, lSPL, MFG, and SFG in both hemispheres, the analysis based on RTs revealed a different set of active brain regions, with only the lPreCUN being common. Hence, this suggests that different neural circuits were activated for subjective confidence evaluation and RT.

We then formulated a model space consisting of two models with plausible forward and backward connections, with the second model incorporating additional direct connections from the rSPL to the lPreCG and lMeFG (Fig. 5A). Similar to the model space for subjective confidence, we performed a BMR on the fully connected model with all the regions and again found a biologically implausible resultant reduced model with lack of connections between parietal and frontal cortices (Jung et al., 2017) and a lower expected probability compared to our current model space. Consequently, we opted for a two-model space that better aligns with our theoretical framework. BMS using the random effects method indicated a preference for the first model in both fast and slow response conditions (Fig. 5B).

**Fig. 5.**
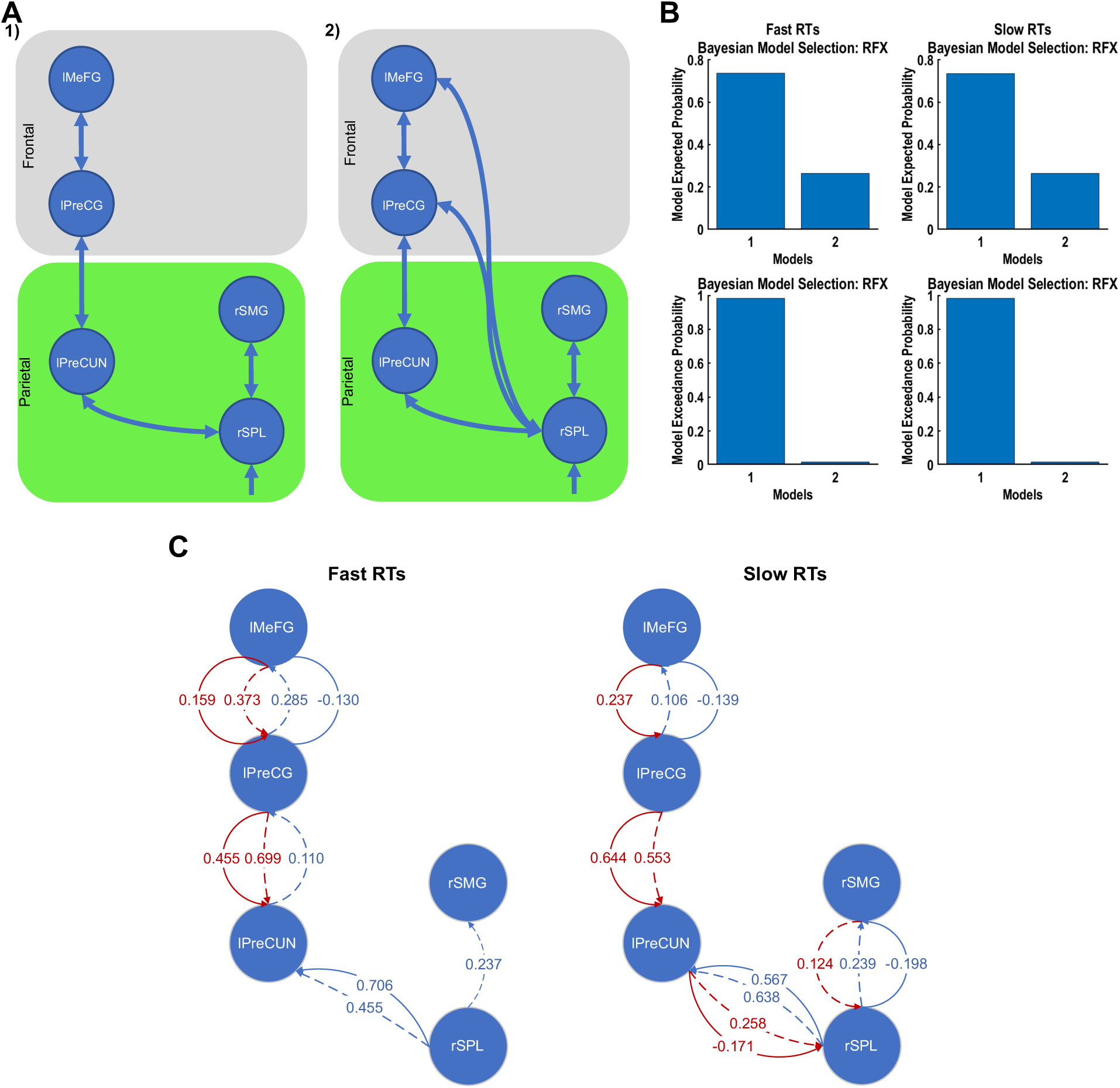
Stronger connections in frontal cortical regions for faster RTs and parietal cortical regions for slower RTs. (**A**) Model space for fast-vs-slow RTs in stimulation phase. Model space for high-vs-low confidence contains plausible connections between active BOLD-fMRI based brain (including parietal (green) and frontal (grey)) regions. (**B**) Bayesian model selection using random-effect approach. Winning model: model 1. (**C**) Forward (blue) and backward (red) connections in the winning model using the PEB approach.

For fast RTs, the winning model exhibited stronger forward connections and more backward connections between the frontal regions (Fig. 5C). However, for slow RTs, there were more backward connections within the parietal cortical regions (Fig. 5C). Compared to slow RTs, the winning model for fast RTs had a stronger SP-to-SS forward connection and a weaker SP-to-DP forward connection between the rSPL and lPreCUN. In addition, the DP-to-SP backward connection between the parietal and frontal regions was weaker, whereas the DP-to-II connection was stronger (Fig. 5C). In addition, for the slow RTs, there were more backward connections in the winning model within the parietal region, suggesting a greater integration of parietal network during slow RTs. The connections in the frontal network follow similar trend as the ones in subjective confidence rating but it is different in the parietal network.

The average MSE of predicted activity against the observed EEG across all the participants for fast (slow) RT condition was 0.0031 (0.0040), with a standard deviation of 0.0015 (0.0023). Visually inspecting the winning model with the best-fitted participant, we observed that the model’s scalp projection activity closely matched the observed scalp EEG activity (Fig. S14). For fast RTs, at 370 ms after stimulus onset, there was negativity in the centroparietal region, which subsequently converted to left hemispheric positivity and right hemispheric negativity at 700 ms. Again, for slow RTs, there was a centroparietal negativity at 370 ms after stimulus onset, which then changed to left hemispheric negativity and right hemispheric positivity at 544 ms. This centroparietal negativity at 370 ms for fast and slow RTs, which subsequently changed to positivity, could be related to the accumulation of sensory evidence towards a decision threshold, as reflected by the CPP observed in EEG signals during perceptual decision-making tasks (O’Connell et al., 2012; Twomey et al., 2015). We observed similar activity only for the low confidence rating condition.

Next, we analysed the source activities for these two RT conditions. The activities of SP and II neural populations in all brain regions were generally higher for slower RTs in the best-fitted participant except for SP activity in rSPL (Fig. 6, dashed), potentially encoding subjective decision uncertainty. In comparison, when averaged across participants, these activities were higher in the rSMG for faster RTs, while other regions showed comparable activities (Fig. 6, solid). The II activities followed a similar pattern to the corresponding SP activities, albeit not as tightly. Thus, averaged source activities were higher at the rSMG for faster RTs, potentially corresponding to an objective measure of high decision confidence. This is also consistent with the higher SP activities observed for higher subjective confidence ratings. We also found involvement of the lPreCUN in encoding response speed as indicated by heightened SS activity during fast RT trials and a trend towards increased DP activity during slow RT trials, though not reaching statistical significance (Fig. S15).

**Fig. 6.**
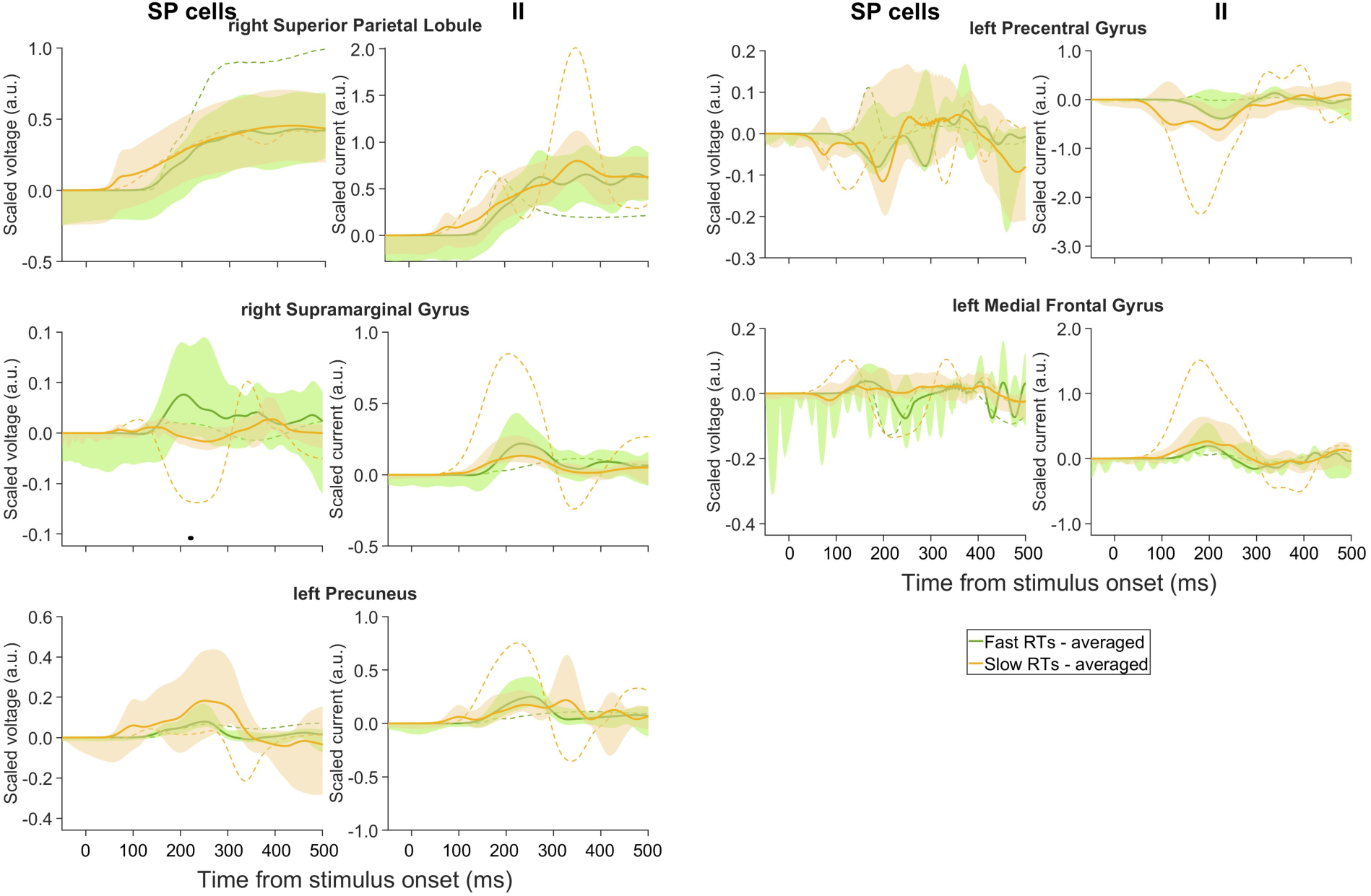
Estimated scaled voltages of SP neural populations and scaled currents of II neural populations of winning model. Green (orange): fast (slow) RTs. Labels as in Fig. 4.

To summarise, generally, there were stronger and more connections in the frontal cortex for faster choice-based RTs while for slower RTs, the parietal cortex had more connections. These trends were similar to that for subjective confidence rating in the frontal cortex although the networks were different.

### Trial-by-trial ERP-DCM predicts parietal cortical layer-based encoding of early decision confidence and uncertainty

We next complement the above standard averaged ERP-DCM approach, in which the lack of considering trial-by-trial variability might potentially mask subtle yet crucial neural dynamics. Specifically, instead of training the DCM on the average ERPs for each participant in each condition, we employed a trial-by-trial ERP-DCM approach using the winning CMC model, identified in the averaged ERP-DCM analysis, to analyse and compare the neural dynamics under different conditions (high versus low confidence ratings and fast versus slow RTs). With this approach, we sought to identify specific and sustained time periods of more than 25 ms (see Figs. S16, S17, S18 and S19 for full results regardless of sustained time periods) in the estimated source activity of the four neural populations in the CMC model for each condition that were significantly correlated with behavioural data among participants using SVR (Drucker et al., 1996) with linear and nonlinear kernels. This was applied in combination with permutation importance (Breiman, 2001), sensitivity analysis (Hinch, 1991), and bootstrapping (Efron, 1979) with a predefined threshold of 70% applied.

First, we found no significant linear correlations between the estimated neural activities and any of the considered four conditions (p < 0.05) and we shall henceforth focus on nonlinear (Gaussian kernel) correlations. With regards to early neural correlates of choice-based RTs, we identified statistically significant correlations over sustained time in the lSPL and lPreCUN regions. Particularly, under high confidence rating condition, estimated SS activities in lSPL exhibited positive correlation with RTs (p < 0.05) for 204-235 ms after stimulus onset, suggesting the encoding of objective decision uncertainty (Fig. 7A). Conversely, under the same condition, estimated DP activities in lPreCUN exhibited significant negative correlation with RTs for 229-269 ms after stimulus onset, suggesting the coding of objective decision confidence in deeper cortical layer (Fig. 7B). Figs. S16 and S17 revealed more detailed results of neural correlation of subjective confidence trials with confidence rating and RT.

**Fig. 7.**
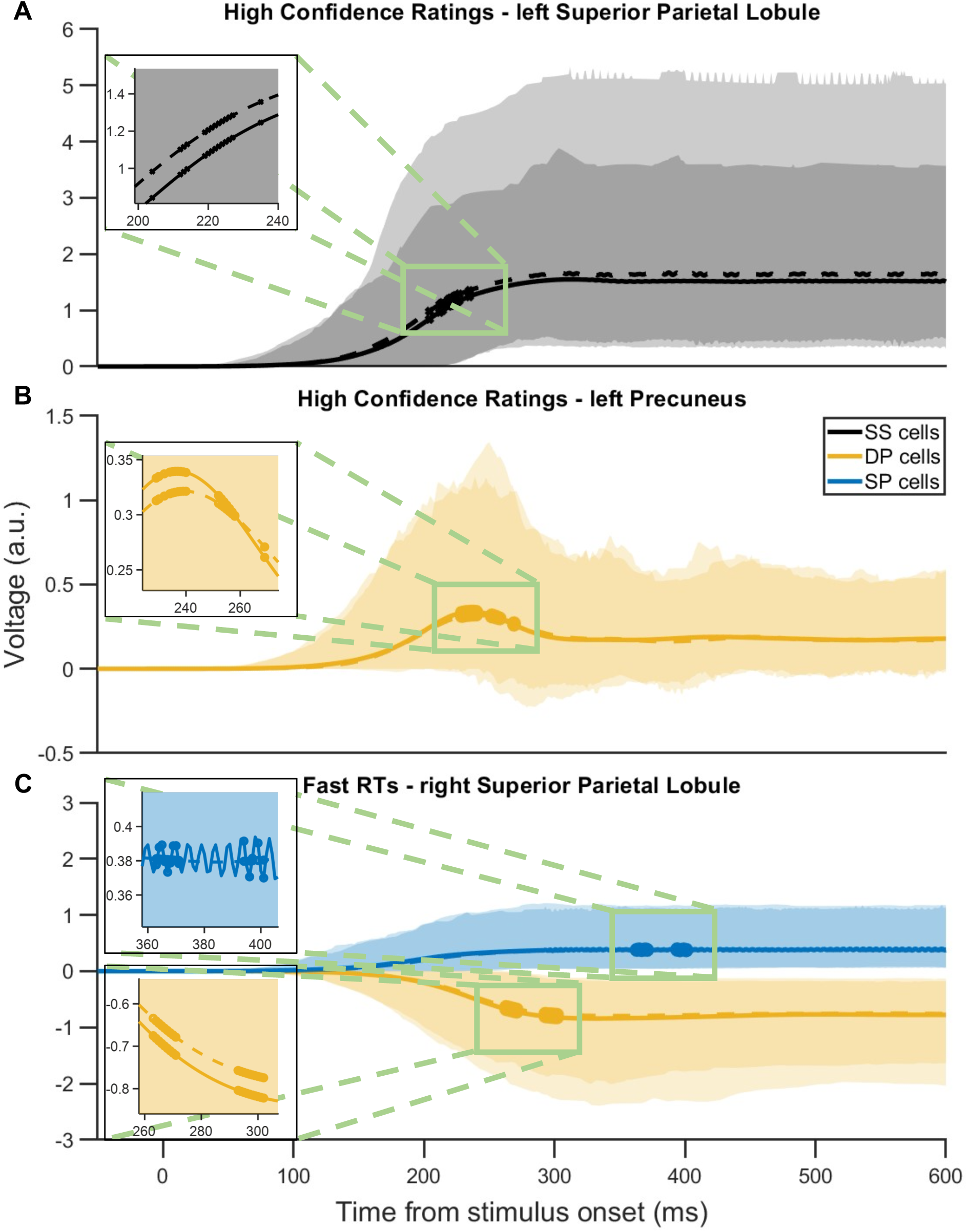
Trial-by-trial DCM identified early decision confidence and uncertainty encoding across parietal cortical layers. (**A-C**) Neural population activities averaged over trials and participants; shaded regions: 95% confidence intervals. (**A-B**) High confidence rating condition. (**A**) Positive correlation between SS activity in lSPL and RTs at 204-235 ms post-stimulus. Solid (dashed) line: fast (slow) RTs. (**B**) Negative correlation between DP activity in lPreCUN and RTs at 229-269 ms post-stimulus. Solid (dashed) line: fast (slow) RTs. (**C**) Fast RT condition. Negative correlation between confidence ratings and SP and DP activities in rSPL at 363-401 ms and 263-302 ms post-stimulus, respectively. Solid (dashed) line: high (low) confidence ratings. Insets: Enlargements at time periods with significant neural activity differences between conditions.

Next for choice-based RT analysis, we focused on trials with fast and slow RTs across all participants, examining the correlation between estimated neural population activities and subjective decision confidence ratings. We found that with fast RTs, there were significant negative correlation between confidence ratings and the estimated SP activities (363-401 ms after stimulus onset) and DP activities (263-302 ms after stimulus onset) in the rSPL (Fig. 7C). This suggests that during these specific time periods with fast decisions, the rSPL might play a role in encoding subjective decision uncertainty. Thus, the winning model predicted rSPL deep-layer neural populations to encode subjective decision confidence. This is to be compared with the above results with lPreCUN’s deep layer encoding objective decision confidence. Figs. S18 and S19 depicted details of all time points of neural activity during RT trials correlated with confidence ratings and RTs.

Overall, our trial-by-trial DCM analysis has illuminated the essential role of the parietal cortex in decision confidence evaluations, revealing neural correlates of both subjective and objective decision confidence/uncertainty in this region early in the decision formation phase, consistent with prior non-human primate study (Kiani & Shadlen, 2009). Specifically, it showed the role of PreCUN in objective confidence evaluation and the role of SPL in the evaluation of both subjective and objective confidence and uncertainty. These results underscore our trial-by-trial ERP-DCM approach in uncovering these nuanced relationships within the brain’s neural circuitry. Importantly, the predicted deep-layer neural population activities do not contribute significantly to scalp EEG data (David et al., 2006). This highlights the potential limitation of solely relying on scalp EEG to understand the neural correlates of decision confidence or uncertainty, as it may not capture crucial information from deeper neural populations.

## Discussion

Previous studies have established a clear relationship between decision confidence levels and the speed in which a decision is made, with faster decisions generally reflecting greater confidence in one’s choice (Pleskac & Busemeyer, 2010; Vickers, 1979). In this work, we delineated the dynamic neural circuit correlate of self-reported subjective decision confidence and that of choice-based RT, which was used as an implicit and more objective measure of decision confidence (Pleskac & Busemeyer, 2010; Vickers, 1979).

To achieve this, we examined the effective connectivity of active brain regions during a perceptual decision-making task with decision confidence assessment, focusing on the early phase of perceptual decision-making. To this end, we employed a combination of ERP and fMRI-informed DCM. We utilised both averaged ERP-DCM and a novel trial-by-trial ERP-DCM approach, contrasting high versus low subjective confidence ratings, as well as fast versus slow choice-based RTs. Our findings revealed distinct frontoparietal networks associated with subjective confidence and RT, despite their significant and negatively correlated behavioural relationship (Figs. 3A and 5A).

When comparing high and low decision confidence ratings, we identified active regions in the parietal cortex (lPreCUN, lIPL, and lSPL) as well as in the frontal cortex (MFG and SFG) (Fig. S2). These findings are consistent with existing literature regarding the role of the parietal cortex in perceptual decision-making and confidence judgments (Cavanna & Trimble, 2006; Kiani & Shadlen, 2009; Platt & Glimcher, 1999; Vickery & Jiang, 2009; Zhou & Freedman, 2019). Specifically, the PreCUN has been implicated in self-referential processing, evaluation of internal states, visuospatial processing, and consciousness (Cavanna & Trimble, 2006). However, the role of the PreCUN in our study contrasts with the findings of <u>Ye et al. (2018)</u> and <u>Zheng et al. (2021)</u>, who demonstrated that the precuneus is causally relevant for mnemonic, but not perceptual, metacognition. This difference suggests that the PreCUN may play a more generalised role in metacognitive processing than previously understood, or that different experimental paradigms reveal distinct aspects of metacognitive function. Similarly, the lSPL has been associated with the integration of sensory information, attentional processing, and decision confidence (Chen et al., 2013), while the lIPL is involved in action planning and prediction (Fogassi & Luppino, 2005). The MFG and SFG, both integral to executive functions and working memory, have also been found to play crucial roles in decision-making processes (Briggs et al., 2021; Rushworth et al., 2004).

When comparing fast and slow RTs, our analysis revealed significant active regions in the parietal cortex (rSPL, lPreCUN, and rSMG) as well as in the frontal cortex (lPreCG and lMeFG). Again, these regions are consistent with their known perceptual decision functions. Specifically, the SMG has been implicated in the integration of sensory information and the evaluation of external stimuli (Davis et al., 2018; Oberhuber et al., 2016) and SPL is associated with attentional control and sensory information integration for decision-making (Molholm et al., 2006).

Our results with the winning DCM did not support direct connections between the (inferior/superior) parietal lobule and the frontal cortex but they were linked via the lPreCUN region. Interestingly, despite the different networks’ involvement in subjective decision confidence evaluation (Fig. 3) and RT (Fig. 5), the lPreCUN was found to be common active brain regions. This shared region may reflect integration point where confidence-related processes could modulate decision-making and RT, aligning with <u>Balsdon et al. (2020)</u> that suggests confidence judgments might influence decision-making. In comparison, the rSPL was active when decisions were fast while lSPL was active with high subjective confidence, possibly reflecting the distinct roles these lateralised regions play in evaluating objective and subjective confidence. These findings indicate that, while distinct, the pathways for confidence and RT may not be entirely independent and could interact at key neural hubs.

By investigating the correlation of estimated neural population activities with subjective decision confidence and choice-based RT in average DCM and trial-by-trial DCM, we found that the winning DCM predicted early neural correlates of subjective decision confidence and choice-based RT in deeper parietal (PreCUN and SPL) cortical layer (Fig. 7). In addition, the model also predicted the parietal (SPL) cortex to be involved in encoding subjective decision uncertainty (Fig. 7C). The results of encoding decision confidence and uncertainty in different regions are consistent with a previous theoretical study which distinguishes the two decision features (Pouget et al., 2016). Our results may not be readily captured by conventional scalp EEG analyses and underscores the importance of investigating neural activity from deeper cortical regions to fully comprehend the intricate circuit mechanisms underpinning decision confidence judgements and decision-making processes (Boucher et al., 2023; Chandrasekaran et al., 2017; Kiani & Shadlen, 2009). Moreover, the best-fitted participant exhibited some neural activity patterns that were vastly different from that of averaged across participants, showing high participant variability (see also the large confidence intervals in Figs. 4 and 6).

To investigate variability between participants, we employed the PEB method on participant-specific effective connectivity parameters retrieved from the winning DCM. This hierarchical Bayesian approach provided participant-level estimated connection strengths and their uncertainty at the group level. In terms of directed functional connectivity, we found, in the winning DCM, stronger forward connections between parietal regions in the low decision confidence rating condition (Fig. 3C). With high subjective confidence, there were stronger forward connections between the lPreCUN and lMFG and more backward connections (Fig. 3C), suggesting a greater integration of the frontoparietal network. This is consistent with the role of the frontoparietal network in integrating sensory information and internal states for decision-making (Heekeren et al., 2008; Summerfield & Tsetsos, 2015). With low subjective confidence, the stronger connections within the parietal cortex may reflect the need for additional sensory integration and attentional processing when confidence is low.

For fast decisions, i.e. fast RTs, stronger forward and more backward connections between the frontal regions were found (Fig. 5C). This is in stark contrast to the winning DCM for high subjective confidence ratings, which exhibited more backward connections in the frontoparietal network. These diverging connectivity patterns suggest that unique neural mechanisms may underpin the evaluation of subjective and objective confidence. Thus, our results suggest that while the frontoparietal network is involved in the early phase of subjective perceptual decision confidence judgements, the specific nature of connectivity within this network may be complex and nuanced.

The findings in this work can inform neural network modelling with explicit neural module(s) that computes decision confidence or uncertainty (Atiya et al., 2019, 2020; Botvinick et al., 2001; Insabato et al., 2010; Jaramillo et al., 2019). In particular, two of these biologically based cortical circuit models (Atiya et al., 2019; Insabato et al., 2010) involved two cortical neural populations with each separately encoding decision confidence, or similar function, and decision uncertainty. Moreover, consistent with our current study that showed stronger parietal cortical connections with higher decision uncertainty, some of these models (Atiya et al., 2019; Botvinick et al., 2001) provided stronger feedback to sensorimotor (e.g. parietal) area when decision uncertainty was higher. However, our findings challenge the assumption in current models that decision confidence and speed share common neural processing resources. Also, as our study suggested that inhibitory interneuron activity might be involved in the encoding of decision confidence/uncertainty, future computational modelling investigations should consider the involvement of inhibitory influences (Atiya et al., 2019; Insabato et al., 2010) as well as assuming different networks for monitoring decision confidence and speed.

One limitation of this study is the exclusion of participants who did not exhibit a significant monotonic correlation between RT and confidence. While this ensured consistency in task engagement, it reduced the number of participants available for analysis. Future studies should aim for larger sample sizes, specifically for the group of participants who do not exhibit a significant correlation between confidence ratings and RTs, to investigate the neural correlates specific to this group. Additionally, due to maintaining a 75% performance rate for each participant without altering the coherence level in the RDM task, this study could not explore network responses to varying evidence qualities. Moreover, the influence of perceived task difficulty, attention, or fatigue on accuracy cannot be fully quantified in this study due to the lack of additional data, such as pupillometry or fatigue measurements. Future studies should incorporate these measures to better assess their impact on performance. The limited number of error trials also constrained our ability to examine the nuances between potential correct and error response networks, despite our analysis efforts (Figs. S6 and S7).

Taken together, our novel integration of fMRI-informed trial-by-trial ERP-DCM represents a significant advancement in the field, providing new and comprehensive insights into the neural circuit dynamics underlying confidence evaluations and decision-making processes. Our findings support the notion that early decision confidence evaluation and decision formation are interrelated processes (Fleming & Daw, 2017; Gherman & Philiastides, 2018; Heereman et al., 2015; Hoven et al., 2022; Zizlsperger et al., 2014), and contribute to the ongoing debate regarding the distinction between these two processes (Kiani & Shadlen, 2009; Pleskac & Busemeyer, 2010; Yeung & Summerfield, 2014). This comprehensive approach paves the way for future research in this domain, offering a valuable framework for investigating the neural correlates of more general decision confidence and uncertainty processing.

## Data and Code Availability

Source codes on subset of processed data (for demonstrative purposes) used in our analysis are available at https://github.com/asadpouretal/DCM-confidence. Original data are available in the open dataset (Gherman & Philiastides, 2020).

## Author Contributions

A.A., K.W.-L. designed and conceptualised analyses. A.A. performed the analysis, modelling and statistical analyses. A.A., K.W.-L. validated the data and analyses. A.A., K.W.-L. wrote the paper. K.W.-L. supervised the research.

## Acknowledgements

We thank Brendan Lenfesty for providing constructive feedback on the manuscript, and Saugat Bhattachayya for helpful discussion on regression method. We are grateful for access to the Tier 2 High Performance Computing resources provided by the Northern Ireland High Performance Computing (NI-HPC) facility funded by the UK Engineering and Physical Sciences Research Council (EPSRC), Grant No. EP/T022175/1.

## Data description

An open concurrent EEG-fMRI dataset on perceptual decision confidence was utilised to investigate the active brain regions and effective connectivity of fMRI-informed EEG data (Gherman & Philiastides, 2020). The dataset comprises 24 participants aged 20–32 years; however, due to inconsistencies in the structural and functional data of the last participant, analyses were conducted on the remaining 23 participants. These participants were right-handed, had normal vision, and no history of neurological disorders (Gherman & Philiastides, 2018).

Participants discriminated the direction of coherent motion in RDM and rated their confidence. The RDM stimuli comprised white dots moving in a black background within a circular aperture, with a subset moving coherently to form the signal, and the rest moving randomly as noise. Task difficulty was controlled by adjusting the proportion of coherently moving dots, with the aim of maintaining overall performance at approximately 75% correct responses, individually calibrated for each participant during a separate training session using a 3-down-1-up staircase procedure. Each trial began with an RDM stimulus presented for a maximum of 1.2 seconds, during which participants made a left or right-directional discrimination using a button press with their right index finger. This was followed by a blank screen and a random delay of 1.5 to 4 s. Subsequently, participants rated their confidence on a white horizontal bar scale for 3 s. Trials ended with another random delay of 1.5 to 4 seconds. Each participant performed two experimental blocks of 160 trials each, corresponding to two separate fMRI runs. All behavioural responses were executed using the right hand on an MR-compatible button box. Fig. 2A shows the sequence within a trial.

A 3-T Siemens MRI scanner was used to record two main task runs, each with 794 brain volumes. Functional data were acquired using a T2*-weighted gradient echo, echo-planar imaging (EPI) sequence with the following parameters: 32 interleaved slices, 0.3 mm gap, 3 × 3 × 3 mm voxel size, 70 × 70 matrix size, 210 mm field of view (FOV), 30 ms echo time (TE), 2000 ms repetition time (TR), and 80° flip angle. Additionally, a high spatial resolution anatomical volume was obtained at the end of the session using a T1-weighted sequence with these parameters: 192 slices, 0.5 mm gap, 1 × 1 × 1 mm voxel size, 256 × 256 matrix size, 256 mm FOV, 2300 ms TE, 2.96 ms TR, and 9° flip angle.

Simultaneously with the fMRI data, EEG data were collected using a 64-channel MR-compatible system from Brain Products, Germany, with the Brain Vision Recorder software at a 5000 Hz sampling rate. Electrodes were positioned according to the 10 – 20 system, with additional nasion, reference, and ground electrodes. In-line resistors ensured participant safety and input impedance was maintained below 25 kΩ. Data acquisition was synchronized with the MRI scanner, and experimental events and participant responses were also recorded and synchronized with the EEG data. For further details, please refer to the original study (Gherman & Philiastides, 2018).

## Data preprocessing

The fMRI data preprocessing procedure applied to the open dataset includes slice-timing correction to adjust for differences in image acquisition times, high-pass filtering with a cutoff of 100 s to remove low-frequency noise, spatial smoothing using a Gaussian kernel of 8 mm to improve signal-to-noise ratio, and head motion correction to account for participants’ movements during the scan. Additionally, we used SPM12 (Penny et al., 2011) for realignment of the fMRI images, extraction of motion parameters, co-registration of the mean fMRI volumes with the structural data for each participant, and normalisation to standard brain space.

In the open dataset, EEG data had been preprocessed using MATLAB. Gradient artefacts were corrected by subtracting average artefact templates from the EEG signal, followed by a 12 ms median filter to remove residual spike artefacts. Standard EEG artefacts were corrected, and a 0.5 – 40 Hz band-pass filter was applied to remove DC drifts and high-frequency noise, with data then downsampled to 1000 Hz. Eye movement and cardiac-related artefacts were minimized using principal component analysis, with data baseline corrected by removing the average signal during the 100 ms prestimulus interval (Gherman & Philiastides, 2018). No additional preprocessing steps were applied in our study.

## Data Analysis

In our study, the data analysis was systematically divided into two pivotal subsections, each crucial for our subsequent DCM analysis. The first subsection, BOLD-fMRI data analysis, was essential in identifying active brain regions and their specific roles in the decision-making process. The second subsection, EEG data analysis, served as a preparatory phase, wherein the EEG data were processed for the DCM analysis. Together, these analyses form the foundation for our DCM analysis.

## BOLD-fMRI data analysis

To statistically analyse the functional data and extract active brain regions under different conditions, we utilised the generalised linear model (GLM) technique in the SPM12 fMRI toolbox (Penny et al., 2011). Specifically, if *Y* is the BOLD response over time for one voxel, the GLM can be formulated as:

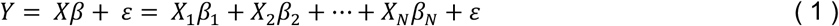

where β is model parameters, *X* is the design matrix for *N* regressors inclusive of effects of interest and no interest, and ε is the residual error vector. The optimal β weights that minimise the squared error values are calculable as (De Martino et al., 2015):

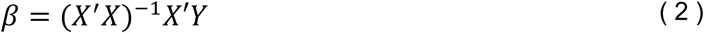

SPM generates the fMRI time series by convolving a time series of delta functions representing the event onsets with HRF, its time derivatives, and factors determining the width of the HRF as basis functions (Penny et al., 2011). Standard HRF utilised extensively in SPM is a mixture of two or more gamma functions (Penny et al., 2011).

For functional data analysis, we employed first-level and second-level statistical analyses to delineate active brain regions under varying conditions. The first-level analysis was conducted within-participant calculations using T-contrast calculations (Penny et al., 2011), a method designed for fMRI signal analysis across multiple conditions, assuming normally distributed data. This step involved calculating the t-statistic for each brain voxel to discern differences between the means of two conditions (high vs. low confidence ratings and fast vs slow RTs). Following this, we adjusted the p-values utilising Bonferroni correction (Bonferroni, 1936) and applied a family-wise error (FWE) correction at a level of 0.05 (Penny et al., 2011). For the second-level analysis, data from all participants were pooled together to facilitate group-level T-contrast analysis.

We conducted GLM analyses across all experimental phases, including stimulation presentation, delay, and confidence rating, to extract active brain regions and compare different conditions within and across these phases (Penny et al., 2011). BOLD signals, inclusive of three condition-specific regressors and six nuisance regressors for motion artefacts, were analysed at three levels: runs, participants, and groups.

During and across all phases, we calculated T-contrasts to compare epochs of high vs. low confidence ratings and fast vs. slow RTs. This identified regions with significant activity differences between these states (Penny et al., 2011). Confidence levels were categorised into high (≥7), medium (5, 6), and low (≤4). RTs were clustered into four groups using k-means clustering with IBM SPSS Statistics (Version 27) (IBM Corp., 2020), computed separately for each participant. This approach allowed us to capture the natural variability in RT distributions without imposing equal trial numbers across bins. Quartile binning was avoided as it would have constrained the groupings into equal-sized bins, potentially misrepresenting the natural variability in RTs. RTs below the second cluster centre were classified as fast, and those above the third cluster centre as slow. We employed T-contrasts to discern active brain regions during each phase, applying family-wise error correction (p < 0.05) to all calculations (Penny et al., 2011). The identified active brain regions will subsequently inform our DCM analysis. Given our interest in early neural correlates of decision confidence, we conducted a further analysis of the stimulation presentation phase.

### EEG data analysis

Using the EEGLAB toolbox (Delorme & Makeig, 2004), we added the events to the dataset. Then we exported it to the SPM12 toolbox for further analysis. Within SPM12, we divided the EEG data into epochs (100 ms before the stimulus onset to 2s after the stimulus onset for each participant based on the onsets of the stimulation phase, and we selected epochs corresponding to various conditions (high and low confidence ratings, fast and slow RTs) during the stimulation phase, and prepared them for DCM implementation. The dynamics from the EEG data will then be used to identify effective connectivity at the source level via DCM, guided by BOLD-fMRI data. Importantly, DCM creates a forward model to map estimated neural activity at the source level to scalp EEG activity using the leadfield matrix, rather than performing source localisation (David et al., 2006; Penny et al., 2011).

### DCM analysis

In our study, we utilised DCM (Penny et al., 2011) to investigate the neural mechanisms underlying early confidence evaluation during perceptual decision-making. DCM enables us to explore directed connectivity between brain regions, thereby providing valuable insights into how these regions interact and contribute to confidence evaluation. To achieve a comprehensive analysis, we employed two types of DCM analyses: averaged ERP-DCM and trial-by-trial ERP-DCM. Averaged ERP-DCM analysis captures general trends and predominant patterns of neural connectivity across all trials, offering a robust overview of the brain’s overall response during the task (David et al., 2006). Furthermore, we implemented an innovative trial-by-trial ERP-DCM approach, delving deeper into the individual trial dynamics. This allows us to dissect the variability and specifics of neural interactions in relation to early confidence evaluation. By integrating these two approaches, we aim to provide a holistic and nuanced understanding of the neural basis of early confidence evaluation in perceptual decision-making.

### Averaged ERP-DCM analysis

In our study, the DCM analysis was instrumental in deciphering the neural mechanisms underlying perceptual confidence and decision-making processes. Our analysis began by meticulously defining the model space, a foundational step that delineates potential connectivity patterns among brain regions based on GLM results (Penny et al., 2011). Insights from the GLM analysis facilitated the formulation of hypotheses regarding possible forward and backward extrinsic connections between active cortical regions.

We were interested in modelling gamma band activities as they have been suggested to be related to perceptual decision-making (Donner et al., 2009). Thus, we selected the CMC neural model, which comprised four distinct neural populations per cortical column, namely SS neural population, II, DP, and SP neural populations (Bastos et al., 2012; Pinotsis et al., 2013). The CMC model is particularly suitable for modelling higher brain frequencies, including gamma activity, due to its inclusion of multiple neural populations within the cortical column, as opposed to conventional ERP-DCMs which may not adequately capture these higher frequencies (Pinotsis et al., 2013). These populations are intricately interconnected, as depicted in Fig. 2B, with each population playing a unique role in modulating neural activity.

For each participant and condition, we averaged the epochs corresponding to high and low confidence ratings, as well as fast and slow choice-based RTs, from the stimulation phase, which spanned from 50 ms prior to stimulus onset to 1.2 seconds after onset. However, our DCMs estimated neural activity until 800 ms post-stimulus onset to particularly investigate early neural correlates.

Neural activity is fundamentally influenced by the fluctuations in voltages and currents across different neural populations. In the CMC model, SP and DP neural populations predominantly affect post-synaptic potentials, serving as vital indicators of neural activity. Excitatory SS neural population contribute to feedforward input, while inhibitory interneurons modulate these dynamics by regulating the currents. A simplified illustration of these relationships is provided here (Fig. 2B), with a comprehensive mathematical depiction available in <u>Pinotsis et al.</u> (Pinotsis et al., 2013).

To probe the neural basis of early confidence evaluation, we employed a cumulative Gaussian signal as our stimulus input, similar to the approach used in <u>FitzGerald et al</u>. (FitzGerald et al., 2015), after also examining a Gaussian bump and sums of two logistic functions. However, the other functions could not estimate the EEG data optimally. The onset of the stimulus was set at approximately 200 ms post-stimulus (Fig. 2C), accounting for the delay in sensory information reaching cortical sensory regions, such as the parietal areas (Lamme & Roelfsema, 2000). Using SPM12, we linearly mapped the post-synaptic voltage, primarily from SP neural population, to scalp EEG activities through a cortical surface patch (Penny et al., 2011). By assigning Gaussian distributions to the parameters and varying extrinsic forward and backward connections (i.e., *A^F^* and *A^B^*) across different DCM models, we estimated the probability density of model parameters, utilising variational free energy and the Laplace approximation method (Penny et al., 2011).

### Statistical analysis of averaged ERP-DCM

After calculating the inverse models in the model space, a BMS identified the winning model using the RFX procedure (Penny et al., 2011). BMS is a statistical method used to compare different models based on their likelihood given the observed data. It is an essential step in model comparison as it informs us about the most probable model that explains the observed data. The RFX approach was chosen because it accounts for the variability between participants, assuming different performance mechanisms for different participants (Penny et al., 2011).

Considering each model’s estimated posterior distribution given the observed data, one can utilise the BMA strategy to calculate the average over all models by averaging the posterior distributions of models weighted by their posterior model probability (Trujillo-Barreto et al., 2004). BMA is a valuable tool as it helps in obtaining a more accurate estimate by considering the uncertainty in model selection.

To investigate variability between participants, we employed PEB on participant-specific effective connectivity parameters retrieved from the winning DCM (Penny et al., 2011). PEB is a hierarchical Bayesian approach that provides participant-level estimated connection strengths and their uncertainty at the group level.

### Trial-by-trial ERP-DCM analysis

In our study, following the averaged ERP-DCM analysis and identification of the winning model for each condition, we employed the trial-by-trial ERP-DCM approach to analyse and compare the neural dynamics of conditions (high vs low confidence rating and fast vs slow RT) during the stimulation phase. We used the connections from the winning model to train the trial-by-trial DCMs with the CMC model.

For each trial, we used the data from the whole epoch, which ranged from -50 ms to 1.2 s from the stimulation onset, to train the DCMs. This range was chosen to ensure that we captured all relevant neural activity during the stimulation phase. The initial parameters of the trial-by-trial DCMs were kept the same as those used in the averaged ERP-DCMs. By employing this innovative approach, we aimed to gain a more comprehensive understanding of the neural basis of early confidence evaluation.

### Statistical analysis of trial-by-trial ERP-DCMs

The statistical analysis of trial-by-trial ERP-DCM was performed to uncover shared neural mechanisms underlying behavioural responses. In our analysis, we were particularly interested in identifying time points in the activity of the four different neural populations in the CMC model—SS neural population, II, DP, and SP neural populations—within the extracted active brain regions (sources) for each condition that have a significant correlation with the behavioural data. We hypothesised that this relationship might exist, as indicated by previous research (Kepecs et al., 2008; Kiani & Shadlen, 2009).

SVR with linear and non-linear (Gaussian) kernels was utilised to establish associations between neural activity patterns and behavioural metrics (Drucker et al., 1996). After selecting the proper window length based on the three lowest RTs among the participants, we trained the SVR models for each participant for each condition, using all of the estimated neural population activities for trials corresponding to each condition. The SVR models were then evaluated for their predictive performance. We then searched for the top 200 important features (time points) within this selected window. These features were then assessed for their importance in predicting behavioural outcomes using permutation importance (Breiman, 2001). This technique involved permuting each feature within the dataset while maintaining the total number of 1250 permutations. We also performed a sensitivity analysis on the form of perturbation to determine if the relationship between the neural population activity and behavioural data (RT or confidence rating) is positive or negative (Hinch, 1991).

The significance of the predictive performances of the SVR models was then scrutinised using bootstrapping (Efron, 1979). This entailed resampling with replacement to construct distributions of predictive performances for each source and neural population. The significance of these performances was determined by calculating confidence intervals through bootstrapping 10,000 times, with a predefined threshold of 70% applied to evaluate the results. The 70% threshold is considered reasonable as it is above the 50% baseline, which is equivalent to random guessing in binary classification (James et al., 2013). Moreover, achieving a performance of 90% or higher might be unrealistic due to the complexity of neural data and substantial noise (Hastie et al., 2009).

After evaluating the performance of SVR models, we conducted sensitivity analysis to determine whether the relationship between neural population activity and behavioural data (RT or confidence rating) is positive or negative. This involved perturbing the data by adding a perturbation factor to the neural activity and re-evaluating the SVR models to observe any changes in the estimated subjective confidence rating or choice-based RTs. Lastly, we identified significantly common features across subjects and conditions using a permutation test with 10,000 permutations (Fisher, 1935). The steps included performing a histogram count of all subjects to extract the data point repeats among subjects, undertaking 10,000 permutations, calculating a threshold for the counts with an alpha level of 0.05, and identifying data points with counts greater than or equal to the threshold. We then selected common features that had a minimum length of 10 ms, regardless of whether they were consecutive or not. Next, we looked for points in the selected common features that covered at least 25 ms, with at most 25 missed points in between, to ensure a robust and meaningful analysis.

### Software and hardware

We employed the EEGLAB toolbox (Delorme & Makeig, 2004) for preprocessing EEG data and utilised the SPM, version 12 (SPM12) toolbox in MATLAB (Version 2022b) for analysing fMRI data, DCM estimations, and statistical analyses For clustering RTs, we utilised IBM SPSS Statistics (Version 27) on a local Windows machine equipped with 14 CPU cores, Intel i9-13900H, and 64GB RAM. All other data analyses were conducted on the Northern Ireland High Performance Computing (NI-HPC) facility using the Kelvin2 system (www.ni-hpc.ac.uk).

