## Supplementary Materials for "Effective connectivity predicts distributed neural coding of perceptual decision confidence, uncertainty and speed"

**This file includes:**

Supplementary Note 1
Table S1
Figs. S1 to S19
References

### Supplementary Note 1

In this Supplementary Information, we provide detailed information and analyses that support the findings reported in the main manuscript. The supplementary materials encompass tables and figures that collectively provide further details on the relationship between decision confidence levels, choice-based reaction times (RTs), and their neural correlates in perceptual decision-making. Below is a summary of each item.

Table S1 lists the fMRI, EEG and EEG-informed fMRI active brain regions that correlate with participants' decision confidence reports during the decision phase based on findings from previous studies, presenting, to date, a comprehensive overview of neural correlates associated with confidence in decision-making tasks. In particular, regions such as the ventral striatum, anterior cingulate cortex (ACC), supplementary motor area (SMA), and multiple frontal and parietal sites have been consistently implicated in decision confidence during stimulus presentation/decision formation phase.

Figures S1 to S19 show the results of our analysis of neural activity response under various conditions:

- **Fig. S1**: Statistical parametric map of the T-statistic (SPM{T}) maps show no significant differences in brain activation between low and high confidence sessions during the stimulation/response phase.
- **Fig. S2**: Significantly higher brain activation differences for high-vs-low confidence rating condition.
- **Fig. S3.** No significant differential brain activation during stimulation/response phase in high-vs-low confidence rating condition when excluding subjects with non-significant RT-confidence relationship. The fMRI SPM{T} analysis parallels

the thresholds and statistical parameters detailed in Fig. S1, with results indicating a lack of significant activation differences.

- **Fig. S4.** No significant differential brain activation during stimulation/response phase in low-vs-high confidence rating condition when excluding subjects with non-significant RT-confidence relationship. The fMRI SPM{T} analysis parallels the thresholds and statistical parameters detailed in Fig. S1, with results indicating a lack of significant activation differences.
- **Fig. S5.** Significant sensory and movement related active regions against implicit baseline during stimulation phase. The fMRI SPM{T} analysis parallels the thresholds and statistical parameters detailed in Fig. S1. Active brain regions are in occipital, parietal, and frontal regions. The extracted areas include undesired regions related to movements, stimulus processing, and other non-relevant brain activities which could obscure the ability to extract key regions specifically relevant to confidence or choice-based RT.
- **Fig. S6**: No significant differential brain activation between correct and incorrect responses during stimulation/response phase. Error trials too low for statistical significance using family-wise error (FWE) correction.
- **Fig. S7**: No significant differential brain activation when comparing incorrect to correct choices during stimulation/response phase. Error trials too low for statistical significance using family-wise error (FWE) correction.
- **Fig. S8**: No significant differential brain activity for fMRI SPM{T} analysis for leftward-vs-rightward stimulus condition during stimulation/response phase.

- **Fig. S9**: No significant differential brain activity for fMRI SPM{T} analysis for rightward-vs-leftward stimulus condition during stimulation/response phase.
- **Fig. S10**: Scalp topography depicting differential EEG activity for high-vs-low confidence rating condition.
- **Fig. S11**: Predominant encoding of confidence rating by DP neural population in left precuneus, while encoding subjective uncertainty by SS neural population in left superior frontal gyrus, illustrating the simultaneous neural encoding of decision confidence and uncertainty.
- **Fig. S12**: Higher brain activation associated with fast-vs-slow choice-based RT condition.
- **Fig. S13**: Significantly higher brain activity in specific regions for slow-vs-fast choice-based RT condition.
- **Fig. S14**: Scalp maps representing observed and model-predicted EEG activity during the stimulation/response phase for trials categorized by choice-based RTs.
- **Fig. S15**: Involvement of left precuneus in encoding response speed, suggesting its potential role in the objective evaluation of decision confidence.
- **Fig. S16**: Correlation of estimated source activity with confidence ratings and choice-based RTs in high confidence rating trials.
- **Fig. S17**: Estimated activities of inhibitory Interneurons (II) in left superior frontal gyrus associated with the encoding of subjective decision uncertainty in low confidence rating trials.
- **Fig. S18**: Estimated SP and DP activities in right superior parietal lobule correlate with subjective decision uncertainty.

- **Fig. S19**: In slow RT trials, left precuneus correlates with confidence rating, while estimated DP activities in right superior parietal lobule correlates with faster choice-based RTs.

**Table S1. Brain region activations correlating with human participants' decision confidence during stimulation/decision formation phase and post-decision phases**. Orange (black): positive (negative) correlation. Abbreviations of brain regions: angular gyrus (AG), inferior parietal gyrus (IPG), calcarine gyrus (CalG), rectal gyrus (RG), ventral striatum (VS), frontal eye field (FEF), medial prefrontal cortex (mPFC), precuneus (PreCUN), medial parietal (mPar), temporal pole (TP), striatum (Str), lateral orbitofrontal cortex (lOFC), anterior prefrontal cortex (aPFC), dorsomedial prefrontal cortex (dmPFC), inferior frontal gyrus (IFG), middle frontal gyrus (MFG), superior frontal gyrus (SFG), dorsolateral prefrontal cortex (dlPFC), ventromedial prefrontal cortex (vmPFC), supplementary eye field (SEF), inferior parietal lobule (IPL), intraparietal sulcus (IPS), middle occipital gyrus (MOG), medial temporal lobe (MTL), superior frontal cortex (SFC), supramarginal gyrus (SMG), anterior insula (aINS), parietal (Par), posterior (Post), cerebellum (Cer), thalamus (Thal), cuneus (Cun), lingual gyrus (Lin), fusiform gyrus (Fus), putamen (Put), medial frontal cortex (MFC), precentral gyrus (PreCG), postcentral gyrus (PostCG), paracentral lobule (ParCL), superior parietal lobule (SPL), basal forebrain (BF), hippocampus (HC), inferior frontal junction (IFJ), inferior temporal gyrus (ITG), superior temporal gyrus (STG), middle temporal gyrus (MTG), superior occipital gyrus (SOG), posterior cingulate cortex (PCC), rostro-lateral prefrontal cortex (rlPFC).

| Publications | Task paradigm | Imaging modality | Task phase | Brain regions correlated with confidence |
|---|---|---|---|---|
| **Heereman et al. (Heereman et al., 2015)** | RDM | fMRI | Stimulation / decision formation | L AG, L IPG, CalG, R Lin, R Fus, vmPFC, RG, L posterior cingulate, SMA/dmPFC, R SFG, L Lin, L Fus, R PreCUN, SPL, R IPL, L IFG |
| **Hebart et al. (Hebart et al., 2016)** | | fMRI | Stimulation / decision formation | VS, ACC, SMA, FEF, IFG, SPL, rMT+, R aINS, R PreCG |
| **Gherman and Philiastides (Gherman & Philiastides, 2018)** | | EEG-informed fMRI | Stimulation / decision formation | Str, lOFC, ACC , Lateral occipital cortex (inferior), MFG (anterior), Occipital pole, R Cer, R ITG, SFG (SMA), dmPFC, R IFG, L PreCG |

| Publications | Task paradigm | Imaging modality | Task phase | Brain regions correlated with confidence |
|---|---|---|---|---|
| **Qiu et al. (Qiu et al., 2018)** | | fMRI | Initial decision | No activation compared to control trials |
| **Li and Yang (Li & Yang, 2012)** | Glass pattern | fMRI | Stimulation / decision formation | R Post MFC, IPS, L SFG, L aINSr, L IFG, L IPL, L Post fusiform |
| **Gherman and Philiastides (Gherman & Philiastides, 2015)** | Face + Object | EEG | Stimulation / decision formation | dmPFC, Par cortex |
| **Qiu et al. (Qiu et al., 2018)** | Sudoku | fMRI | Initial decision | IFJ |
| **Jaeger et al. (Jaeger et al., 2020)** | Gap Location | fMRI | Stimulation / decision formation | BF, R SEFs, IPL, SMG, MFG, Cer, MOG, Put, visual Thal, and right PostCG |
| **Shapiro and Grafton (Shapiro & Grafton, 2020)** | Approach–avoidance task | fMRI | Stimulation / decision formation | STG, MTG, L Cun, SOG, R Lin, R SMG, R PostCG, R Inferior Par, R SMG, R Inferior Par, R MFG, R Median cingulate and paracingulate |

| Publications | Task paradigm | Imaging modality | Task phase | Brain regions correlated with confidence |
|---|---|---|---|---|
| | | | | gyri, L Anterior cingulate and paracingulate gyri, R SFG, dorsolateral, R MFG, R PreCUN, R MFG |
| **Hoven et al. (Hoven et al., 2022)** | Gabor patches | fMRI | Stimulation / decision formation | vmPFC, PCC, dlPFC, rlPFC, aINS, R Put, R IFG, SMA, mid-ACC, ACC, IPL |
| **Post-decision brain activations** | | | | |
| **Pereira et al. (Pereira et al., 2020)** | Box with dots | EEG-informed fMRI | Delay period between post-decision formation and confidence rating | Occipital, VS, L Put, L vmPFC, R HC, SMA, dACC, L SFC, L MFC, IFG, IPL, L aPFC, L MTL , L PreCUN |
| **Bang and Fleming (Bang & Fleming, 2018)** | RDM | fMRI | Unspecified | perigenual ACC |
| **Fleming et al. (Fleming et al., 2018)** | | fMRI | Post-decision confidence rating | medial aPFC, PreCUN/ mPar, Temporal lobe, White matter, L Inf. Par, R Sup. Occipital, |

| Publications | Task paradigm | Imaging modality | Task phase | Brain regions correlated with confidence |
|---|---|---|---|---|
| | | | | MFG, R PreCG, ParCL, R Cer, Occipital / inf. Par, Thal, pMFC, lateral aPFC |
| **Hilgenstock et al. (Hilgenstock et al., 2014)** | Grating Orientation Task | fMRI | Delay period between post-decision formation and confidence rating | superior medial gyrus bilateral, R aPFC, R dlPFC |
| **Morales et al. (Morales et al., 2018)** | Word-Shape recognition task | fMRI | Confidence/follow rating | left PreCG, left PostCG, Post midline VS, vmPFC, dACC/pre-SMA, Par cortex, bilateral PFC |

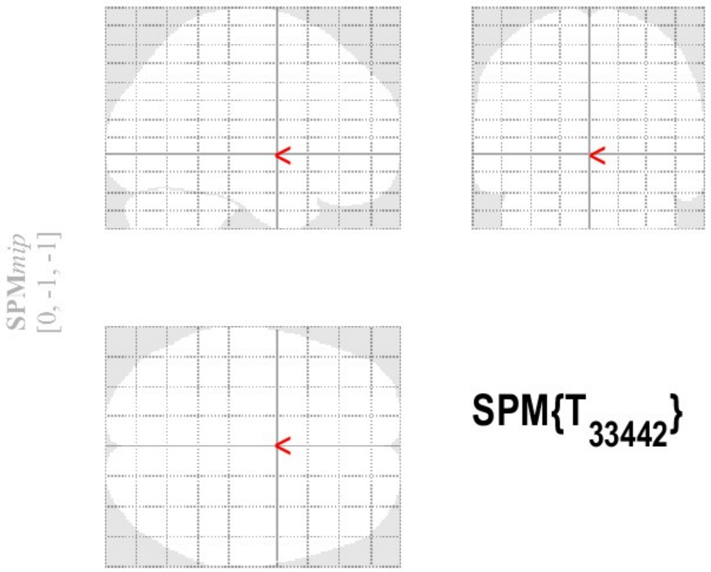


SPM{$T_{33442}$}

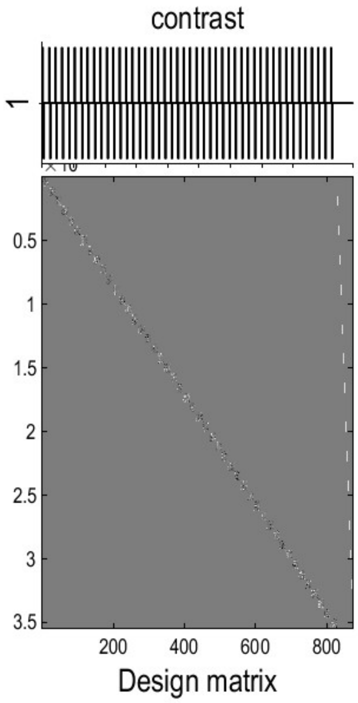


**SPMresults:** .\Second Level Stats
Height threshold T = 4.471160 {p<0.05 (FWE)}
Extent threshold k = 0 voxels

**Statistics:** ***p-values adjusted for search volume***

| set-level | | cluster-level | | | | peak-level | | | | | mm mm mm |
|---|---|---|---|---|---|---|---|---|---|---|---|
| $p$ | $c$ | $p_{FWE-corr}$ | $q_{FDR-corr}$ | $k_E$ | $p_{uncorr}$ | $p_{FWE-corr}$ | $q_{FDR-corr}$ | $T$ | $(Z_E)$ | $p_{uncorr}$ | |

***no suprathreshold clusters***

*table shows 3 local maxima more than 8.0mm apart*

Height threshold: T = 4.47, p = 0.000 (0.050)
Extent threshold: k = 0 voxels
Expected voxels per cluster, <k> = 2.836
Expected number of clusters, <c> = 0.05
FWEp: 4.471, FDRp: Inf, FWEc: Inf, FDRc: Inf

Degrees of freedom = [1.0, 33442.0]
FWHM = 12.3 12.5 13.0 mm mm mm; 4.1 4.2 4.3 {voxels}
Volume: 939924 = 34812 voxels = 404.6 resels
Voxel size: 3.0 3.0 3.0 mm mm mm; (resel = 73.84 voxels)

**Fig. S1. Statistical parametric mapping (SPM) analysis demonstrating non-significant brain region activation.** The SPM{T} maps depict the lack of suprathreshold clusters when contrasting low confidence against high confidence sessions, across all sessions during the stimulation/response phase. Thresholds were set at a height of $T = 4.471160$, $p < 0.05$ (family-wise error corrected), and an extent threshold of $k = 0$ voxels, indicating no significant activation. The design matrix and contrast vector are provided, alongside the adjusted p-values for search volume, which confirm the absence of significant differences. The table below the SPM{T} maps specify that there are no local maxima surpassing the 8.0 mm separation threshold, further supporting the null findings.

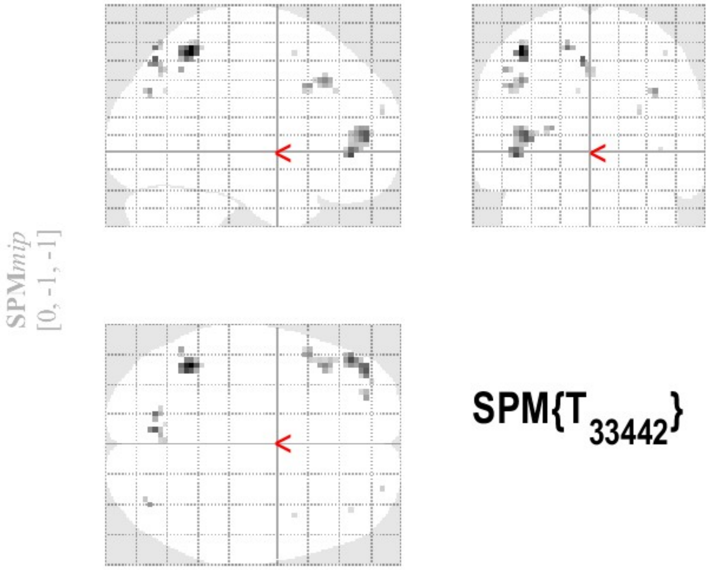


SPM{$T_{33442}$}

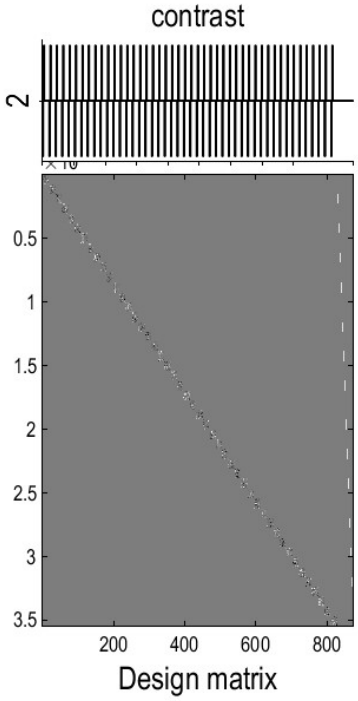


**SPMresults:** .\Second Level Stats
Height threshold T = 4.471160 {p<0.05 (FWE)}
Extent threshold k = 0 voxels

### Statistics: *p-values adjusted for search volume*

| set-level | | cluster-level | | | | peak-level | | | | | mm mm mm | | |
|---|---|---|---|---|---|---|---|---|---|---|---|---|---|
| $p$ | $c$ | $p_{FWE-corr}$ | $q_{FDR-corr}$ | $k_E$ | $p_{uncorr}$ | $p_{FWE-corr}$ | $q_{FDR-corr}$ | $T$ | $(Z_E)$ | $p_{uncorr}$ | | | |
| 0.000 | 12 | 0.000 | 0.030 | 26 | 0.005 | 0.000 | 0.110 | 5.53 | 5.53 | 0.000 | -42 | -52 | 56 |
| | | 0.000 | 0.007 | 43 | 0.001 | 0.003 | 0.254 | 5.15 | 5.15 | 0.000 | -45 | 41 | -4 |
| | | | | | | 0.003 | 0.254 | 5.09 | 5.09 | 0.000 | -39 | 50 | 8 |
| | | 0.004 | 0.220 | 9 | 0.073 | 0.003 | 0.254 | 5.09 | 5.08 | 0.000 | -6 | -73 | 50 |
| | | 0.009 | 0.411 | 5 | 0.171 | 0.005 | 0.321 | 4.99 | 4.99 | 0.000 | -15 | -73 | 56 |
| | | 0.000 | 0.035 | 22 | 0.009 | 0.009 | 0.438 | 4.87 | 4.87 | 0.000 | -42 | 26 | 38 |
| | | | | | | 0.011 | 0.438 | 4.83 | 4.83 | 0.000 | -48 | 17 | 35 |
| | | 0.015 | 0.489 | 3 | 0.285 | 0.012 | 0.438 | 4.81 | 4.81 | 0.000 | 36 | -76 | 32 |
| | | 0.015 | 0.489 | 3 | 0.285 | 0.015 | 0.497 | 4.76 | 4.76 | 0.000 | -24 | 50 | 11 |
| | | 0.019 | 0.512 | 2 | 0.384 | 0.017 | 0.497 | 4.73 | 4.73 | 0.000 | -51 | -58 | 44 |
| | | 0.019 | 0.512 | 2 | 0.384 | 0.037 | 0.911 | 4.55 | 4.55 | 0.000 | 27 | 59 | 20 |
| | | 0.028 | 0.547 | 1 | 0.547 | 0.044 | 0.969 | 4.51 | 4.51 | 0.000 | 42 | 8 | 53 |
| | | 0.028 | 0.547 | 1 | 0.547 | 0.048 | 0.969 | 4.48 | 4.48 | 0.000 | -9 | -67 | 32 |
| | | 0.028 | 0.547 | 1 | 0.547 | 0.048 | 0.969 | 4.48 | 4.48 | 0.000 | 39 | 41 | -1 |

*table shows 3 local maxima more than 8.0mm apart*

Height threshold: T = 4.47, p = 0.000 (0.050)
Extent threshold: k = 0 voxels
Expected voxels per cluster, <k> = 2.836
Expected number of clusters, <c> = 0.05
FWEp: 4.471, FDRp: Inf, FWEc: 1, FDRc: 22

Degrees of freedom = [1.0, 33442.0]
FWHM = 12.3 12.5 13.0 mm mm mm; 4.1 4.2 4.3 {voxels}
Volume: 939924 = 34812 voxels = 404.6 resels
Voxel size: 3.0 3.0 3.0 mm mm mm; (resel = 73.84 voxels)

**Fig. S2. Significant brain activation during stimulation/response phase in high-vs-low confidence rating condition.** SPM results illustrate marked activation in the IPL (left hemisphere), MFG (both left and right hemispheres), PreCUN (left hemisphere), SPL (left hemisphere), Angular Gyrus (right hemisphere), and SFG (both left and right hemispheres). These findings are indicative of varied neural involvement correlating with confidence levels during task performance when high confidence sessions are more active than low confidence sessions. The fMRI SPM{T} analysis parallels the thresholds and statistical parameters detailed in Fig. S1. The accompanying results table confirms the presence of three distinct local maxima separated by more than 8.0 mm, validating the significant clusters identified.

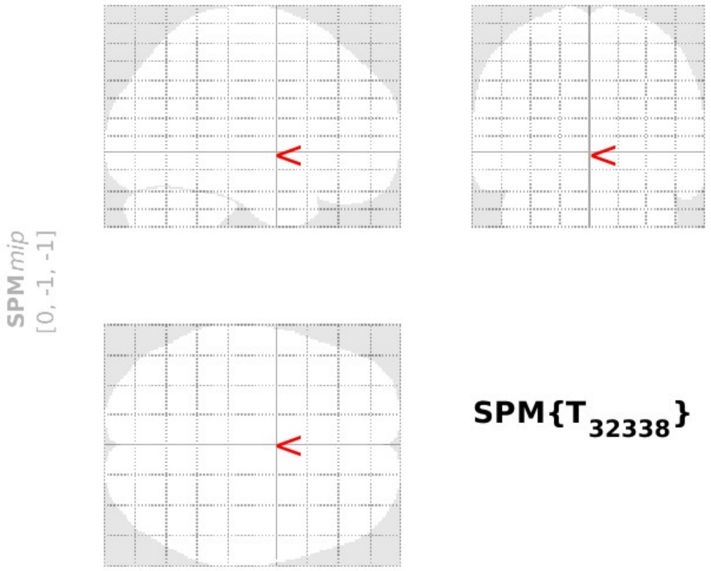


SPM{$T_{32338}$}

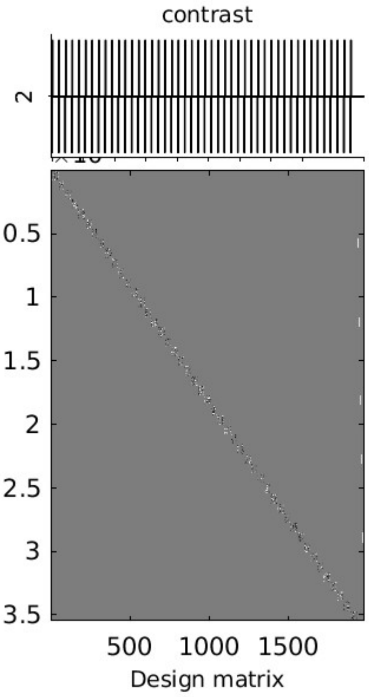


**SPMresults:** ..igh conditions in all phases
Height threshold T = 4.475290 {p<0.05 (FWE)}
Extent threshold k = 0 voxels

**Statistics:** ***p-values adjusted for search volume***

| set-level | | cluster-level | | | | peak-level | | | | | mm mm mm |
|---|---|---|---|---|---|---|---|---|---|---|---|
| $p$ | $c$ | $p_{FWE-corr}$ | $q_{FDR-corr}$ | $k_E$ | $p_{uncorr}$ | $p_{FWE-corr}$ | $q_{FDR-corr}$ | $T$ | $(Z_E)$ | $p_{uncorr}$ | |

***no suprathreshold clusters***

*table shows 3 local maxima more than 8.0mm apart*

Height threshold: T = 4.48, p = 0.000 (0.050)
Extent threshold: k = 0 voxels
Expected voxels per cluster, <k> = 2.779
Expected number of clusters, <c> = 0.05
FWEp: 4.475, FDRp: Inf, FWEc: Inf, FDRc: Inf

Degrees of freedom = [1.0, 32338.0]
FWHM = 12.2 12.4 12.9 mm mm mm; 4.1 4.1 4.3 {voxels}
Volume: 939924 = 34812 voxels = 411.8 resels
Voxel size: 3.0 3.0 3.0 mm mm mm; (resel = 72.54 voxels)

**Fig. S3. No significant differential brain activation during stimulation/response phase in high-vs-low confidence rating condition when excluding subjects with non-significant RT-confidence relationship.** The fMRI SPM{T} analysis parallels the thresholds and statistical parameters detailed in Fig. S1, with results indicating a lack of significant activation differences.

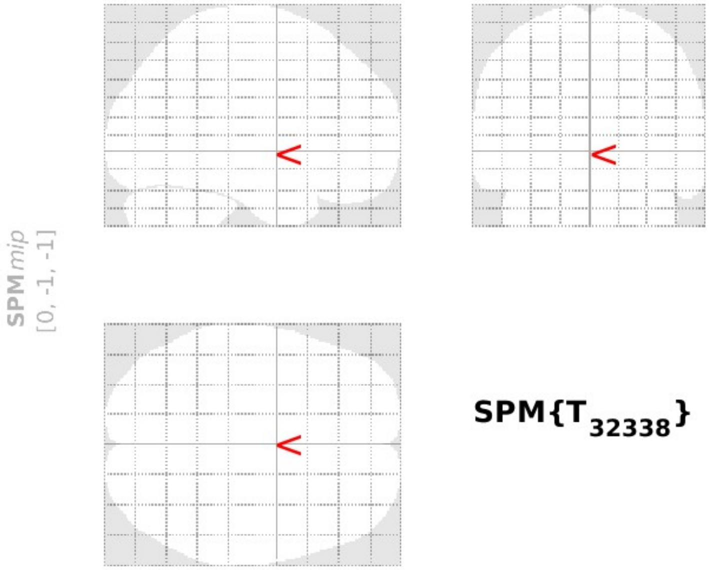


SPM{$T_{32338}$}

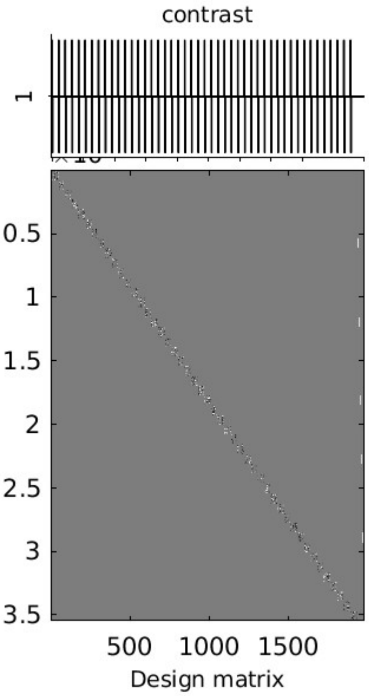


**SPMresults:** ..igh conditions in all phases
Height threshold T = 4.475290 {p<0.05 (FWE)}
Extent threshold k = 0 voxels

**Statistics:** ***p-values adjusted for search volume***

| set-level | | cluster-level | | | | peak-level | | | | | mm mm mm |
|---|---|---|---|---|---|---|---|---|---|---|---|
| $p$ | $c$ | $p_{FWE-corr}$ | $q_{FDR-corr}$ | $k_E$ | $p_{uncorr}$ | $p_{FWE-corr}$ | $q_{FDR-corr}$ | $T$ | $(Z_E)$ | $p_{uncorr}$ | |

***no suprathreshold clusters***

*table shows 3 local maxima more than 8.0mm apart*

Height threshold: T = 4.48, p = 0.000 (0.050)
Extent threshold: k = 0 voxels
Expected voxels per cluster, <k> = 2.779
Expected number of clusters, <c> = 0.05
FWEp: 4.475, FDRp: Inf, FWEc: Inf, FDRc: Inf

Degrees of freedom = [1.0, 32338.0]
FWHM = 12.2 12.4 12.9 mm mm mm; 4.1 4.1 4.3 {voxels}
Volume: 939924 = 34812 voxels = 411.8 resels
Voxel size: 3.0 3.0 3.0 mm mm mm; (resel = 72.54 voxels)

**Fig. S4. No significant differential brain activation during stimulation/response phase in low-vs-high confidence rating condition when excluding subjects with non-significant RT-confidence relationship.** The fMRI SPM{T} analysis parallels the thresholds and statistical parameters detailed in Fig. S1, with results indicating a lack of significant activation differences.

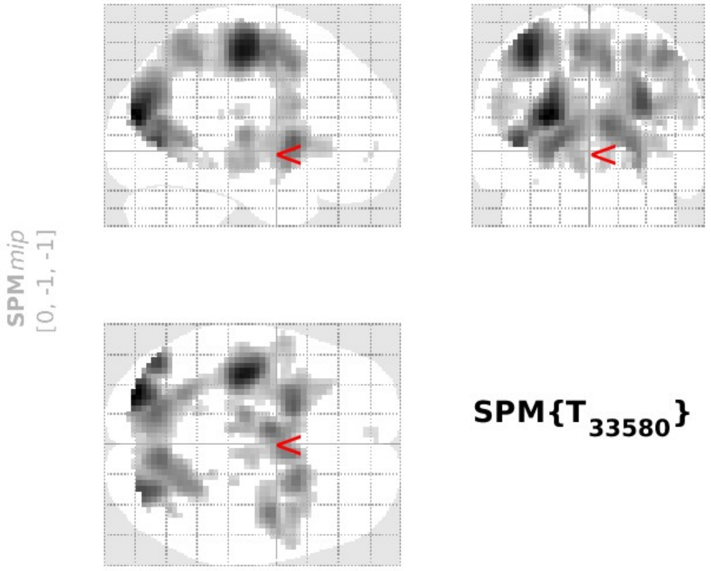


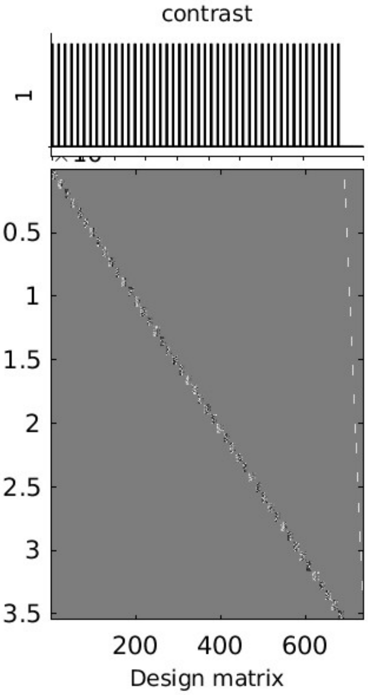


**SPMresults:** .. Stats for stimulation phase
Height threshold T = 4.474987 {p<0.05 (FWE)}
Extent threshold k = 0 voxels

**Statistics:** ***p-values adjusted for search volume***

| set-level | | cluster-level | | | | peak-level | | | | | mm mm mm |
|---|---|---|---|---|---|---|---|---|---|---|---|
| $p$ | $c$ | $p_{FWE-corr}$ | $q_{FDR-corr}$ | $k_E$ | $p_{uncorr}$ | $p_{FWE-corr}$ | $q_{FDR-corr}$ | $T$ | $(Z_E)$ | $p_{uncorr}$ | |
| **0.000** | **13** | **0.000** | **0.000** | **2796** | **0.000** | **0.000** | **0.000** | **14.39** | **Inf** | **0.000** | **-24 -85 20** |
| | | | | | | 0.000 | 0.000 | 13.20 | Inf | 0.000 | -36 -22 56 |
| | | | | | | 0.000 | 0.000 | 11.12 | Inf | 0.000 | -45 -70 2 |
| | | **0.000** | **0.000** | **402** | **0.000** | **0.000** | **0.000** | **10.15** | **Inf** | **0.000** | **-21 8 2** |
| | | | | | | 0.000 | 0.001 | 6.01 | 6.01 | 0.000 | -30 26 -1 |
| | | **0.000** | **0.000** | **498** | **0.000** | **0.000** | **0.000** | **9.62** | **Inf** | **0.000** | **-6 -4 56** |
| | | | | | | 0.000 | 0.000 | 8.76 | Inf | 0.000 | 9 8 47 |
| | | | | | | 0.000 | 0.000 | 8.63 | Inf | 0.000 | 6 2 53 |
| | | **0.000** | **0.000** | **242** | **0.000** | **0.000** | **0.000** | **9.15** | **Inf** | **0.000** | **21 11 2** |
| | | | | | | 0.000 | 0.000 | 8.41 | Inf | 0.000 | 27 11 -4 |
| | | | | | | 0.000 | 0.000 | 6.12 | 6.12 | 0.000 | 36 14 2 |
| | | **0.000** | **0.000** | **415** | **0.000** | **0.000** | **0.000** | **8.88** | **Inf** | **0.000** | **42 -7 50** |
| | | | | | | 0.000 | 0.000 | 7.17 | 7.16 | 0.000 | 39 8 23 |
| | | | | | | 0.000 | 0.001 | 6.02 | 6.02 | 0.000 | 48 2 38 |
| | | **0.000** | **0.000** | **212** | **0.000** | **0.000** | **0.000** | **7.95** | **Inf** | **0.000** | **-15 -19 8** |
| | | | | | | 0.000 | 0.000 | 6.75 | 6.74 | 0.000 | -9 -19 -7 |
| | | | | | | 0.001 | 0.030 | 5.29 | 5.29 | 0.000 | -21 -28 -4 |
| | | **0.000** | **0.000** | **113** | **0.000** | **0.000** | **0.000** | **6.15** | **6.15** | **0.000** | **9 -16 -7** |
| | | | | | | 0.000 | 0.000 | 6.13 | 6.13 | 0.000 | 9 -16 5 |
| | | **0.001** | **0.021** | **19** | **0.013** | **0.000** | **0.010** | **5.52** | **5.52** | **0.000** | **-51 -25 20** |
| | | **0.005** | **0.139** | **7** | **0.107** | **0.001** | **0.022** | **5.36** | **5.36** | **0.000** | **24 -25 -7** |
| | | **0.019** | **0.411** | **2** | **0.379** | **0.010** | **0.217** | **4.86** | **4.86** | **0.000** | **-12 -25 38** |
| | | **0.014** | **0.331** | **3** | **0.280** | **0.014** | **0.284** | **4.79** | **4.78** | **0.000** | **-36 -34 17** |
| | | **0.004** | **0.125** | **8** | **0.087** | **0.018** | **0.368** | **4.72** | **4.72** | **0.000** | **-6 53 -4** |
| | | **0.027** | **0.543** | **1** | **0.543** | **0.039** | **0.786** | **4.53** | **4.53** | **0.000** | **-33 -4 -19** |

*table shows 3 local maxima more than 8.0mm apart*

Height threshold: T = 4.47, p = 0.000 (0.050)
Extent threshold: k = 0 voxels
Expected voxels per cluster, <k> = 2.782
Expected number of clusters, <c> = 0.05
FWEp: 4.475, FDRp: 5.292, FWEc: 1, FDRc: 19

Degrees of freedom = [1.0, 33580.0]
FWHM = 12.2 12.4 12.9 mm mm mm; 4.1 4.1 4.3 {voxels}
Volume: 939924 = 34812 voxels = 411.4 resels
Voxel size: 3.0 3.0 3.0 mm mm mm; (resel = 72.62 voxels)

**Fig. S5. Significant sensory and movement related active regions against implicit baseline during stimulation phase.** The fMRI SPM{T} analysis parallels the thresholds and statistical parameters detailed in Fig. S1. Active brain regions are in occipital, parietal, and frontal regions. The extracted areas include undesired regions related to movements, stimulus processing, and other non-relevant brain activities which could obscure the ability to extract key regions specifically relevant to confidence or choice-based RT.

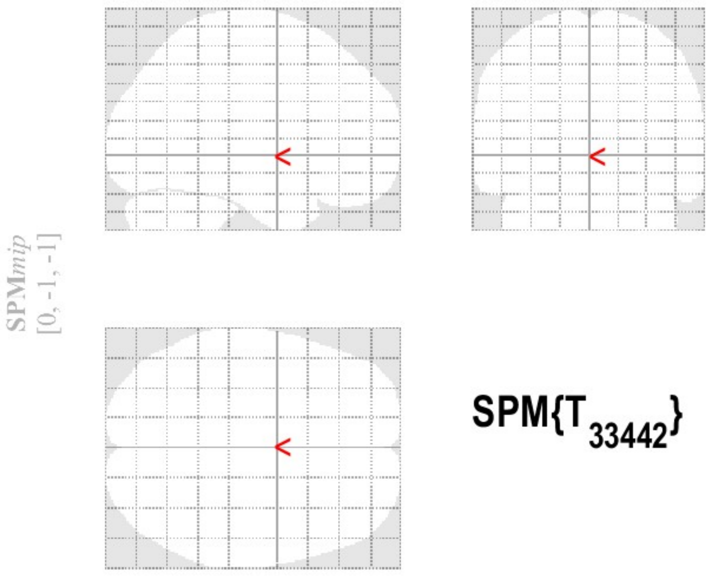


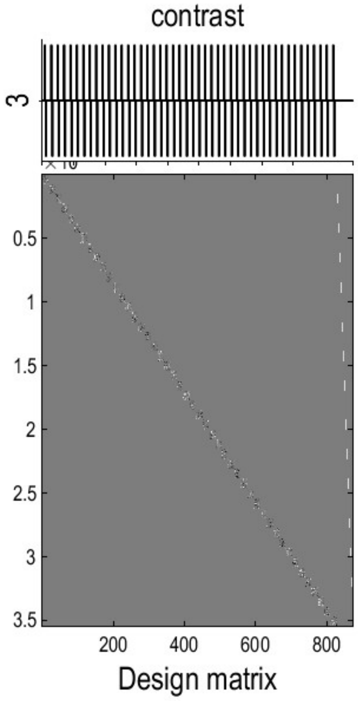


**SPMresults:** .\Second Level Stats
Height threshold T = 4.471160 {p<0.05 (FWE)}
Extent threshold k = 0 voxels

**Statistics:** ***p-values adjusted for search volume***

| set-level | | cluster-level | | | | peak-level | | | | | mm mm mm |
|---|---|---|---|---|---|---|---|---|---|---|---|
| $p$ | $c$ | $p_{FWE-corr}$ | $q_{FDR-corr}$ | $k_E$ | $p_{uncorr}$ | $p_{FWE-corr}$ | $q_{FDR-corr}$ | $T$ | $(Z_E)$ | $p_{uncorr}$ | |

***no suprathreshold clusters***

*table shows 3 local maxima more than 8.0mm apart*

Height threshold: T = 4.47, p = 0.000 (0.050)
Extent threshold: k = 0 voxels
Expected voxels per cluster, <k> = 2.836
Expected number of clusters, <c> = 0.05
FWEp: 4.471, FDRp: Inf, FWEc: Inf, FDRc: Inf

Degrees of freedom = [1.0, 33442.0]
FWHM = 12.3 12.5 13.0 mm mm mm; 4.1 4.2 4.3 {voxels}
Volume: 939924 = 34812 voxels = 404.6 resels
Voxel size: 3.0 3.0 3.0 mm mm mm; (resel = 73.84 voxels)

**Fig. S6. No significant differential brain activation between correct and incorrect responses during the stimulation/response phase.** The fMRI SPM{T} analysis parallels the thresholds and statistical parameters detailed in Fig. S1, with results indicating a lack of significant activation differences. The data affirm the consistency of brain activity irrespective of response accuracy.

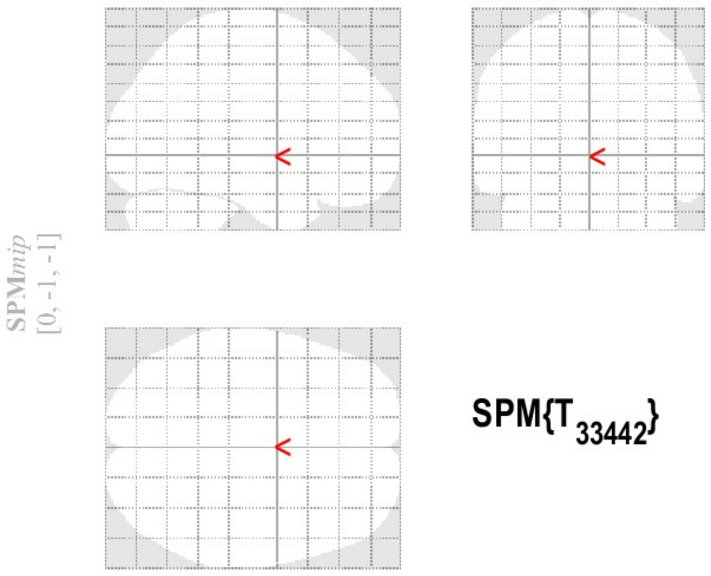


SPM{$T_{33442}$}

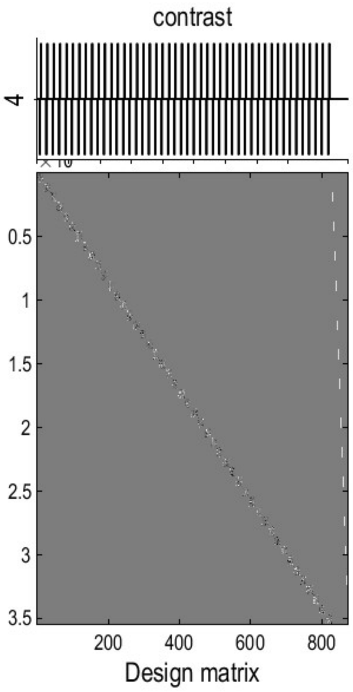


**SPMresults:** .\Second Level Stats
Height threshold T = 4.471160 {p<0.05 (FWE)}
Extent threshold k = 0 voxels

**Statistics:** ***p-values adjusted for search volume***

| set-level | | cluster-level | | | | peak-level | | | | | mm mm mm |
|---|---|---|---|---|---|---|---|---|---|---|---|
| *p* | *c* | $p_{FWE-corr}$ | $q_{FDR-corr}$ | $k_E$ | $p_{uncorr}$ | $p_{FWE-corr}$ | $q_{FDR-corr}$ | *T* | $(Z_E)$ | $p_{uncorr}$ | |

***no suprathreshold clusters***

*table shows 3 local maxima more than 8.0mm apart*

Height threshold: T = 4.47, p = 0.000 (0.050)
Extent threshold: k = 0 voxels
Expected voxels per cluster, <k> = 2.836
Expected number of clusters, <c> = 0.05
FWEp: 4.471, FDRp: Inf, FWEc: Inf, FDRc: Inf

Degrees of freedom = [1.0, 33442.0]
FWHM = 12.3 12.5 13.0 mm mm mm; 4.1 4.2 4.3 {voxels}
Volume: 939924 = 34812 voxels = 404.6 resels
Voxel size: 3.0 3.0 3.0 mm mm mm; (resel = 73.84 voxels)

**Fig. S7. No significant differential brain activation when comparing incorrect to correct choices during the stimulation/response phase.** This fMRI SPM{T} analysis adheres to the thresholds and statistical parameters specified in Fig. S1's caption, and similarly, it reveals no significant activation differences.

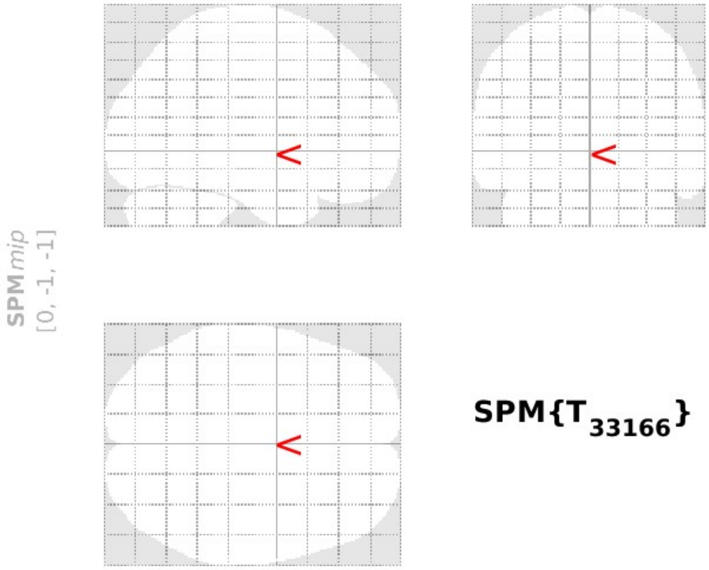


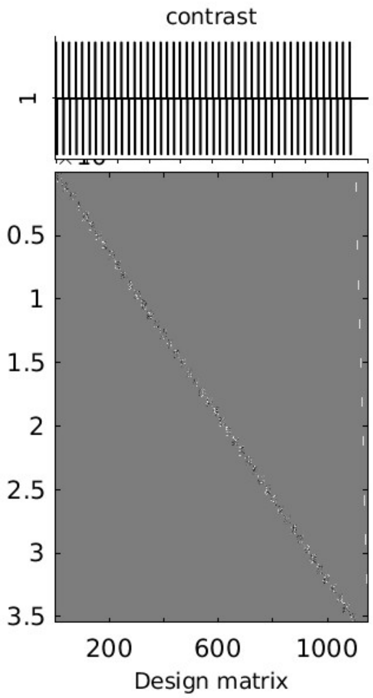


**SPMresults:** ..ect conditions in all phases
Height threshold T = 4.471866 {p<0.05 (FWE)}
Extent threshold k = 0 voxels

**Statistics:** ***p-values adjusted for search volume***

| set-level | | cluster-level | | | | peak-level | | | | | mm mm mm |
|---|---|---|---|---|---|---|---|---|---|---|---|
| $p$ | $c$ | $p_{FWE-corr}$ | $q_{FDR-corr}$ | $k_E$ | $p_{uncorr}$ | $p_{FWE-corr}$ | $q_{FDR-corr}$ | $T$ | $(Z_E)$ | $p_{uncorr}$ | |

***no suprathreshold clusters***

*table shows 3 local maxima more than 8.0mm apart*

Height threshold: T = 4.47, p = 0.000 (0.050)
Extent threshold: k = 0 voxels
Expected voxels per cluster, <k> = 2.826
Expected number of clusters, <c> = 0.05
FWEp: 4.472, FDRp: Inf, FWEc: Inf, FDRc: Inf

Degrees of freedom = [1.0, 33166.0]
FWHM = 12.3 12.5 13.0 mm mm mm; 4.1 4.2 4.3 {voxels}
Volume: 939924 = 34812 voxels = 405.8 resels
Voxel size: 3.0 3.0 3.0 mm mm mm; (resel = 73.62 voxels)

**Fig. S8. No significant differential brain activity for fMRI SPM{T} analysis for leftward-vs-rightward stimulus condition during stimulation/response phase.** This indicates a lack of differential brain activity between the directions of stimuli. For statistical thresholds and parameters, refer to Fig. S1. The results suggest that the directionality of stimulus during the stimulation phase does not elicit a distinct neural response pattern.

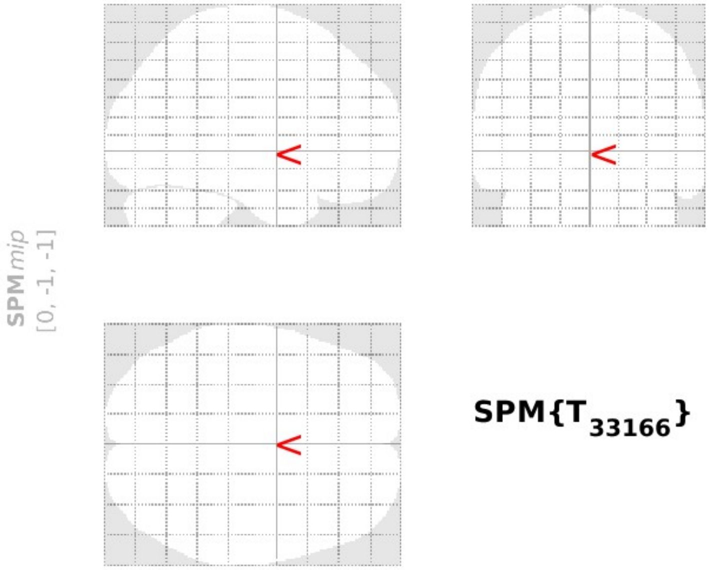


**SPM{$T_{33166}$}**

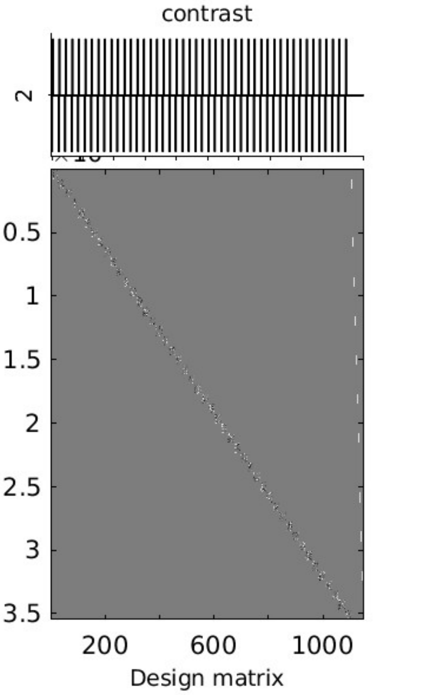


**SPMresults:** ..ect conditions in all phases
Height threshold T = 4.471866 {p<0.05 (FWE)}
Extent threshold k = 0 voxels

**Statistics:** ***p-values adjusted for search volume***

| set-level | | cluster-level | | | | peak-level | | | | | mm mm mm |
|---|---|---|---|---|---|---|---|---|---|---|---|
| $p$ | $c$ | $p_{FWE-corr}$ | $q_{FDR-corr}$ | $k_E$ | $p_{uncorr}$ | $p_{FWE-corr}$ | $q_{FDR-corr}$ | $T$ | $(Z_E)$ | $p_{uncorr}$ | |

***no suprathreshold clusters***

*table shows 3 local maxima more than 8.0mm apart*

Height threshold: T = 4.47, p = 0.000 (0.050)
Extent threshold: k = 0 voxels
Expected voxels per cluster, <k> = 2.826
Expected number of clusters, <c> = 0.05
FWEp: 4.472, FDRp: Inf, FWEc: Inf, FDRc: Inf

Degrees of freedom = [1.0, 33166.0]
FWHM = 12.3 12.5 13.0 mm mm mm; 4.1 4.2 4.3 {voxels}
Volume: 939924 = 34812 voxels = 405.8 resels
Voxel size: 3.0 3.0 3.0 mm mm mm; (resel = 73.62 voxels)

**Fig. S9. No significant differential brain activity for fMRI SPM{T} analysis for rightward-vs-leftward stimulus condition during stimulation/response phase.** No suprathreshold clusters were observed, suggesting similar neural engagement for stimuli directionality. Refer to the statistical details in Fig. S1 for comparison. These findings indicate that the direction of stimulus in the stimulation phase is not a differentiating factor for neural activation.

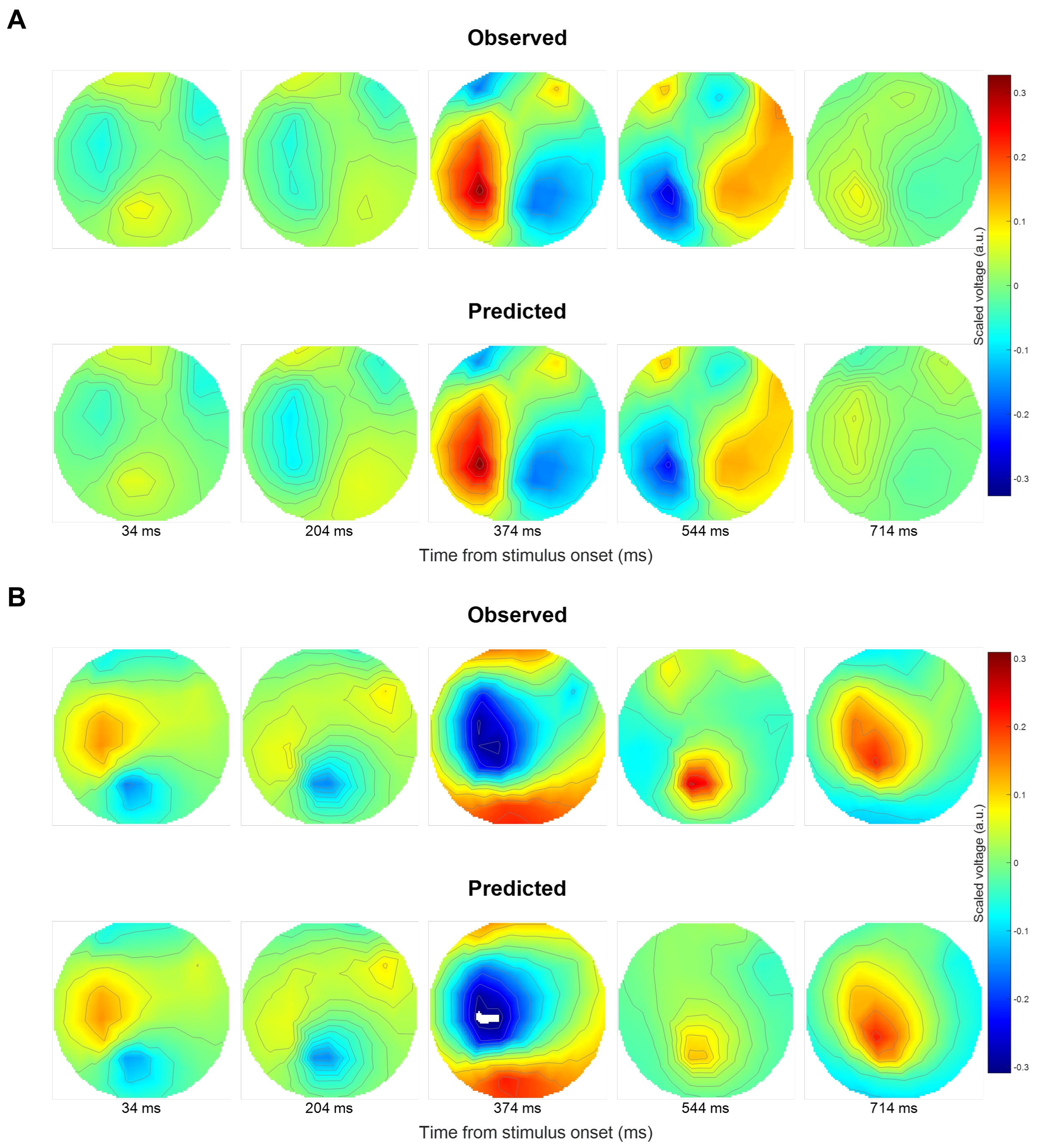


**Fig. S10. Scalp topography depicting differential EEG activity patterns for high versus low confidence trials in the most accurately modelled participant, as per the fMRI-informed EEG-DCM analysis**. (A) EEG scalp activity for high confidence trials, where the winning fMRI-informed EEG- Dynamic causal modelling (DCM) model for the best-fitted participant indicates a left-hemispheric positivity and right-hemispheric negativity at 374 ms post-stimulus. This activity pattern then reverses into left-hemisphere negativity and right-hemisphere positivity by 544 ms. (B) low confidence trials, with centro-parietal negativity at 370 ms evolving into a widespread positivity by 714 ms. These maps align closely with recorded EEG scalp distributions, validating the predictive capability of the EEG-DCM model in capturing the spatial-temporal dynamics of neural activity associated with confidence levels.

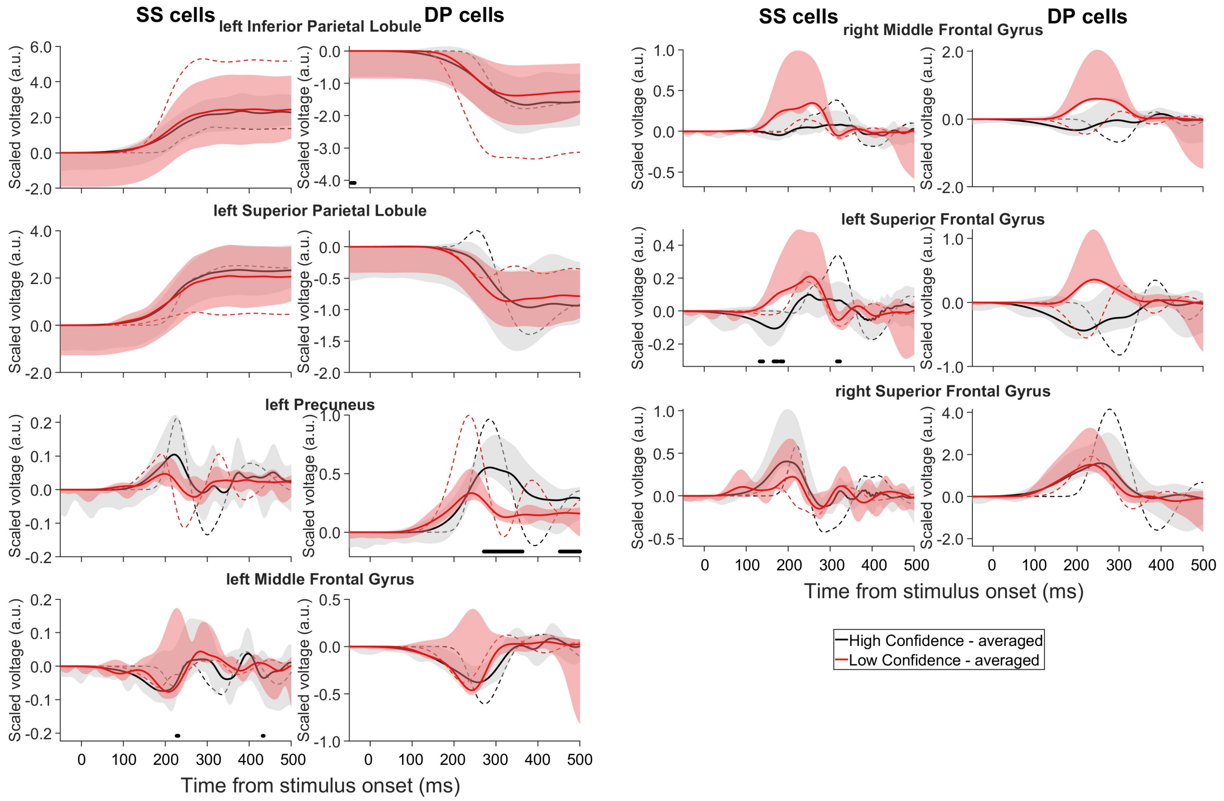


**Fig. S11. Predominant encoding of confidence rating by DP activity in left PreCUN and potential encoding of subjective uncertainty by SS activity in left SFG.** Estimated scaled voltage for SS neural population and scaled current for DP neural populations from the model that was deemed most accurate. Black (red) lines: High (low) confidence ratings. Solid (dashed) lines: averages across participants (best-fitted participant). Shaded areas: 95% confidence. Filled markers above horizontal axis: time points where significant differences were found between the averaged conditions ($p < 0.05$). Left SFG shows increased SS activity during low confidence trials, suggesting encoding of decision uncertainty, whereas left PreCUN exhibits greater DP activity during high confidence trials, indicating encoding of subjective confidence.

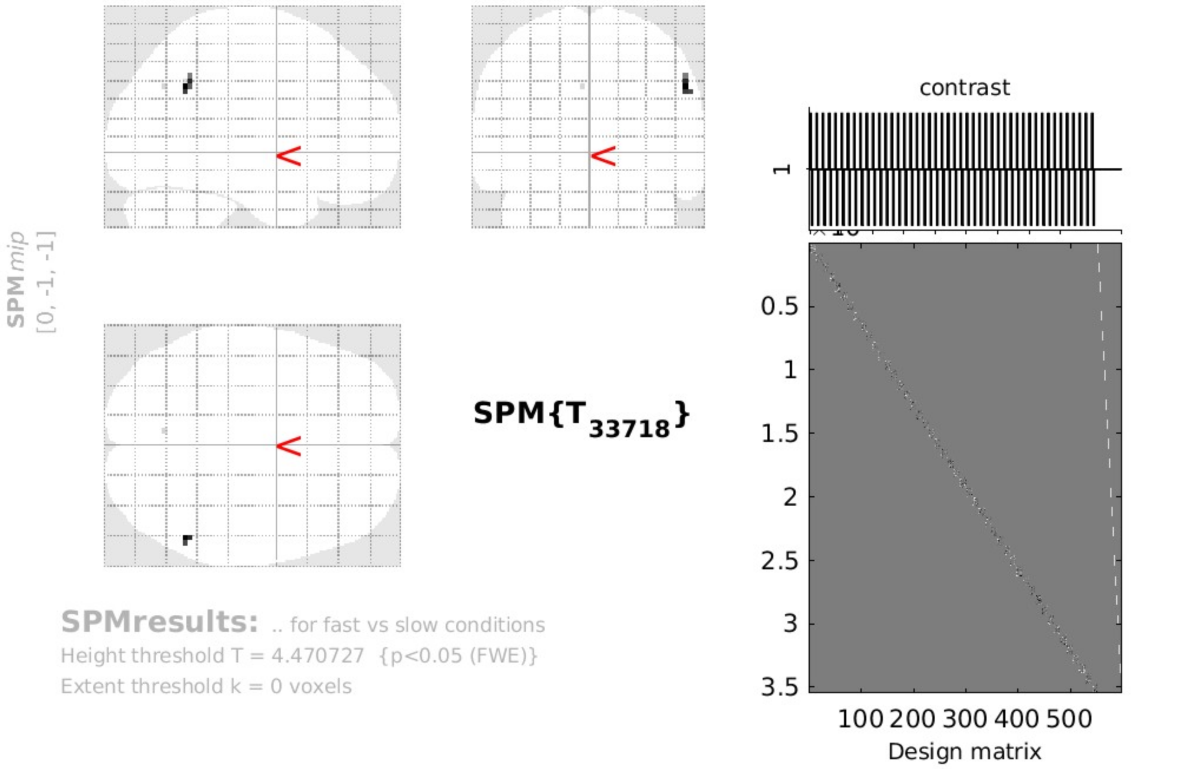


**Statistics:** ***p-values adjusted for search volume***

| set-level | | cluster-level | | | | peak-level | | | | | |
|---|---|---|---|---|---|---|---|---|---|---|---|
| $p$ | $c$ | $p_{FWE-corr}$ | $q_{FDR-corr}$ | $k_E$ | $p_{uncorr}$ | $p_{FWE-corr}$ | $q_{FDR-corr}$ | $T$ | $(Z_E)$ | $p_{uncorr}$ | mm mm mm |
| 0.001 | 2 | 0.006 | 0.221 | 7 | 0.110 | 0.014 | 0.566 | 4.77 | 4.77 | 0.000 | 54 -55 35 |
| | | 0.028 | 0.547 | 1 | 0.547 | 0.042 | 0.834 | 4.52 | 4.51 | 0.000 | -6 -67 35 |

*table shows 3 local maxima more than 8.0mm apart*

Height threshold: T = 4.47, p = 0.000 (0.050)
Extent threshold: k = 0 voxels
Expected voxels per cluster, <k> = 2.843
Expected number of clusters, <c> = 0.05
FWEp: 4.471, FDRp: Inf, FWEc: 1, FDRc: Inf

Degrees of freedom = [1.0, 33718.0]
FWHM = 12.3 12.5 13.0 mm mm mm; 4.1 4.2 4.3 {voxels}
Volume: 939924 = 34812 voxels = 403.8 resels
Voxel size: 3.0 3.0 3.0 mm mm mm; (resel = 73.98 voxels)

**Fig. S12. Higher brain activation associated with fast-vs-slow choice-based RT condition.** SMG in right parietal lobe and the PreCUN in left parietal lobe were significantly more active during fast RTs compared to slow RTs. This pattern of activation correlates with the efficiency of response execution. For detailed statistical thresholds and parameters, see Fig. S1 caption.

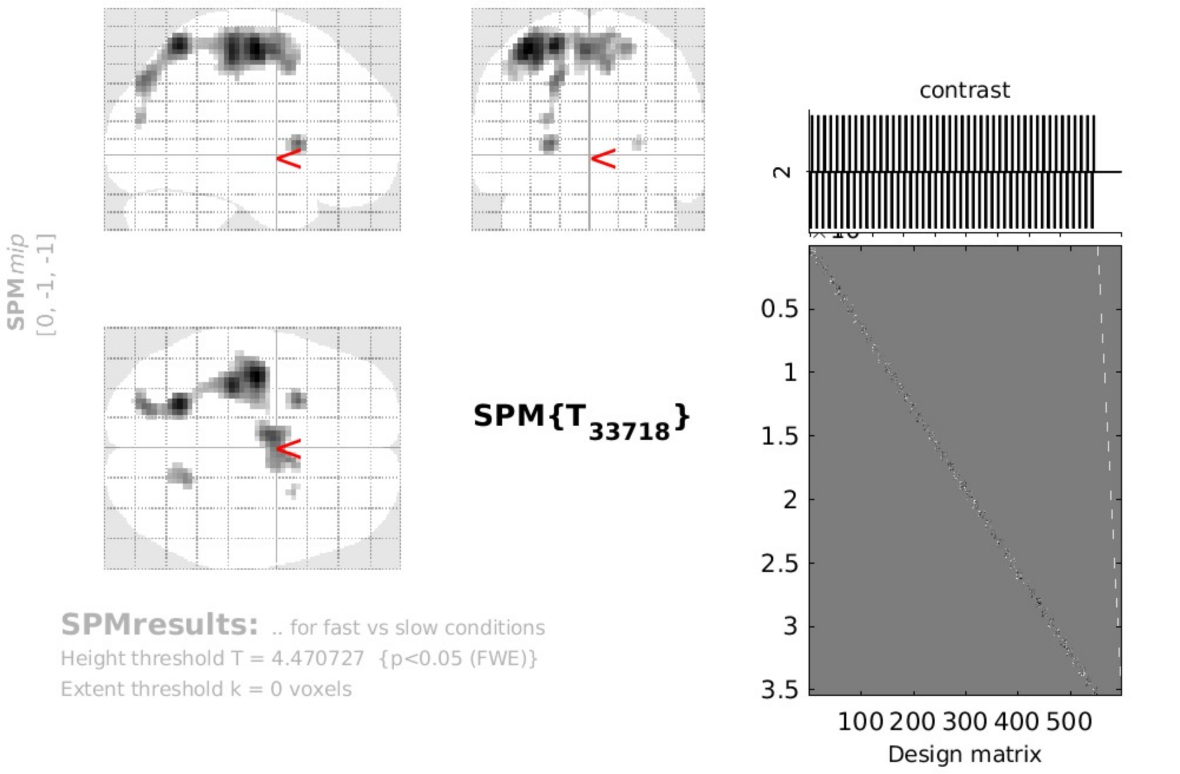


**Statistics:** ***p-values adjusted for search volume***

| set-level | | cluster-level | | | | peak-level | | | | | mm mm mm |
|---|---|---|---|---|---|---|---|---|---|---|---|
| $p$ | $c$ | $p_{FWE-corr}$ | $q_{FDR-corr}$ | $k_E$ | $p_{uncorr}$ | $p_{FWE-corr}$ | $q_{FDR-corr}$ | $T$ | $(Z_E)$ | $p_{uncorr}$ | |
| **0.000** | **5** | **0.000** | **0.000** | **638** | **0.000** | **0.000** | **0.000** | **7.87** | **Inf** | **0.000** | **-39 -13 56** |
| | | | | | | 0.000 | 0.000 | 7.48 | 7.48 | 0.000 | -21 -58 59 |
| | | | | | | 0.000 | 0.000 | 7.33 | 7.33 | 0.000 | -33 -28 56 |
| | | **0.000** | **0.000** | **238** | **0.000** | **0.000** | **0.000** | **6.66** | **6.66** | **0.000** | **-3 -4 56** |
| | | | | | | 0.000 | 0.002 | 5.88 | 5.88 | 0.000 | 6 2 53 |
| | | **0.000** | **0.001** | **39** | **0.001** | **0.000** | **0.000** | **6.33** | **6.33** | **0.000** | **-24 11 5** |
| | | **0.000** | **0.000** | **53** | **0.000** | **0.000** | **0.003** | **5.77** | **5.77** | **0.000** | **18 -55 59** |
| | | **0.004** | **0.074** | **9** | **0.074** | **0.004** | **0.070** | **5.08** | **5.08** | **0.000** | **27 8 5** |

*table shows 3 local maxima more than 8.0mm apart*

Height threshold: T = 4.47, p = 0.000 (0.050)
Extent threshold: k = 0 voxels
Expected voxels per cluster, <k> = 2.843
Expected number of clusters, <c> = 0.05
FWEp: 4.471, FDRp: 5.466, FWEc: 9, FDRc: 39

Degrees of freedom = [1.0, 33718.0]
FWHM = 12.3 12.5 13.0 mm mm mm; 4.1 4.2 4.3 {voxels}
Volume: 939924 = 34812 voxels = 403.8 resels
Voxel size: 3.0 3.0 3.0 mm mm mm; (resel = 73.98 voxels)

**Fig. S13. Significantly higher brain activity in specific regions for slow-vs-fast choice-based RT condition.** The regions include left PreCG, left medial frontal gyrus, left lentiform nucleus (sub-lobar), right SPL, and right extra-nuclear area (sub-lobar). Refer to Fig. S1 for statistical thresholds and parameters. This pattern suggests that slower RTs are associated with increased neural activities in areas implicated in motor control, cognitive processing, and sensory integration.

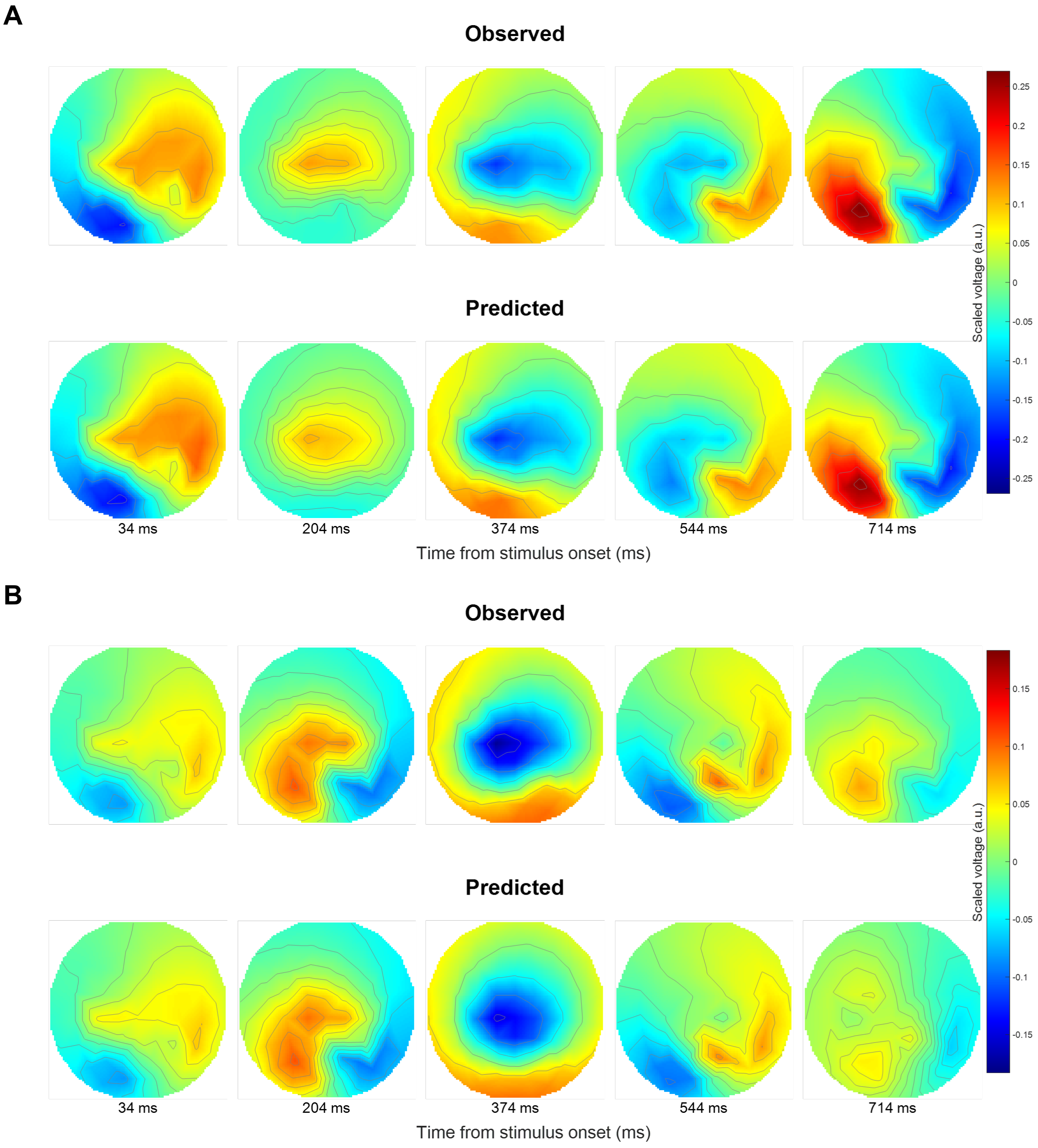


**Fig. S14. Scalp maps representing observed and model-predicted EEG activity during the stimulation/response phase for trials categorized by choice-based RTs**. (A) Trials with fast RT, showing a centro-parietal negativity at 370 ms post-stimulus onset, transitioning to a positivity in the left hemisphere and negativity in the right hemisphere at 700 ms. (B) Trials with slow RT, where initial centro-parietal negativity at 370 ms is followed by a left-hemispheric negativity and a right-hemispheric positivity by 544 ms. These patterns indicate a clear distinction in neural processing speed and lateralization associated with the speed of participants' responses. The consistency of topographical distribution between predicted and observed EEG activity underscores the precision of the EEG-DCM model in capturing the temporal dynamics of neural responses related to RTs.

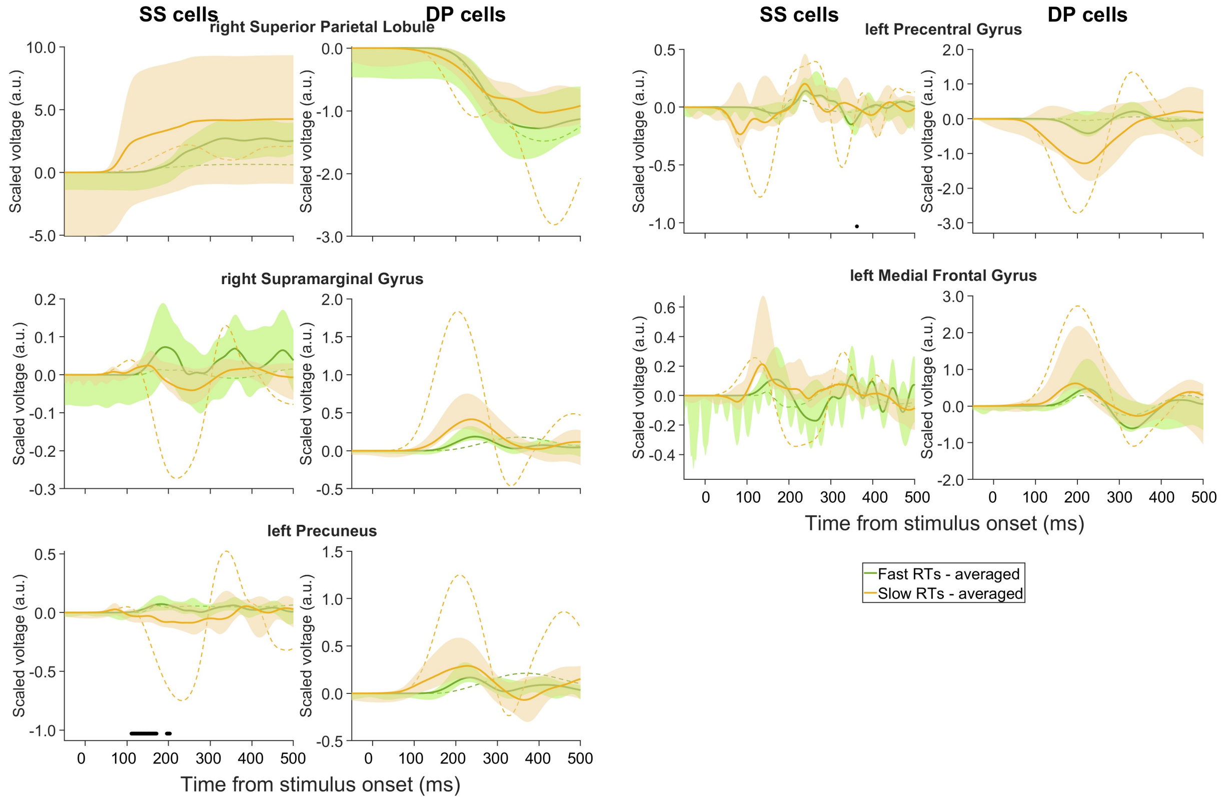


**Fig. S15. Involvement of left PreCUN in encoding fast RT, suggesting objective evaluation of decision confidence.** Estimated scaled voltage for SS neural populations and scaled current for DP neural populations in the winning model, with fast choice-based RTs (green) and slow choice-based RTs (orange). Across most regions, SS activity is consistent between slow and fast choice-based RT trials, except in the left PreCUN, where there is a marked increase in activity during fast trials, underscoring its potential role in the objective evaluation of confidence. In comparison, DP activity is generally higher for slow RT trials in various brain regions across participants, although these differences are not statistically significant. Solid (dashed) lines: averages across participants (best-fitted participant). Shaded areas: 95% confidence. Filled markers above horizontal axis: time points where significant differences were found between the averaged conditions ($p < 0.05$). Left PreCUN shows increased SS activity during fast RT trials, suggesting objective evaluation of decision confidence whereas no significant DP activity difference was observed.

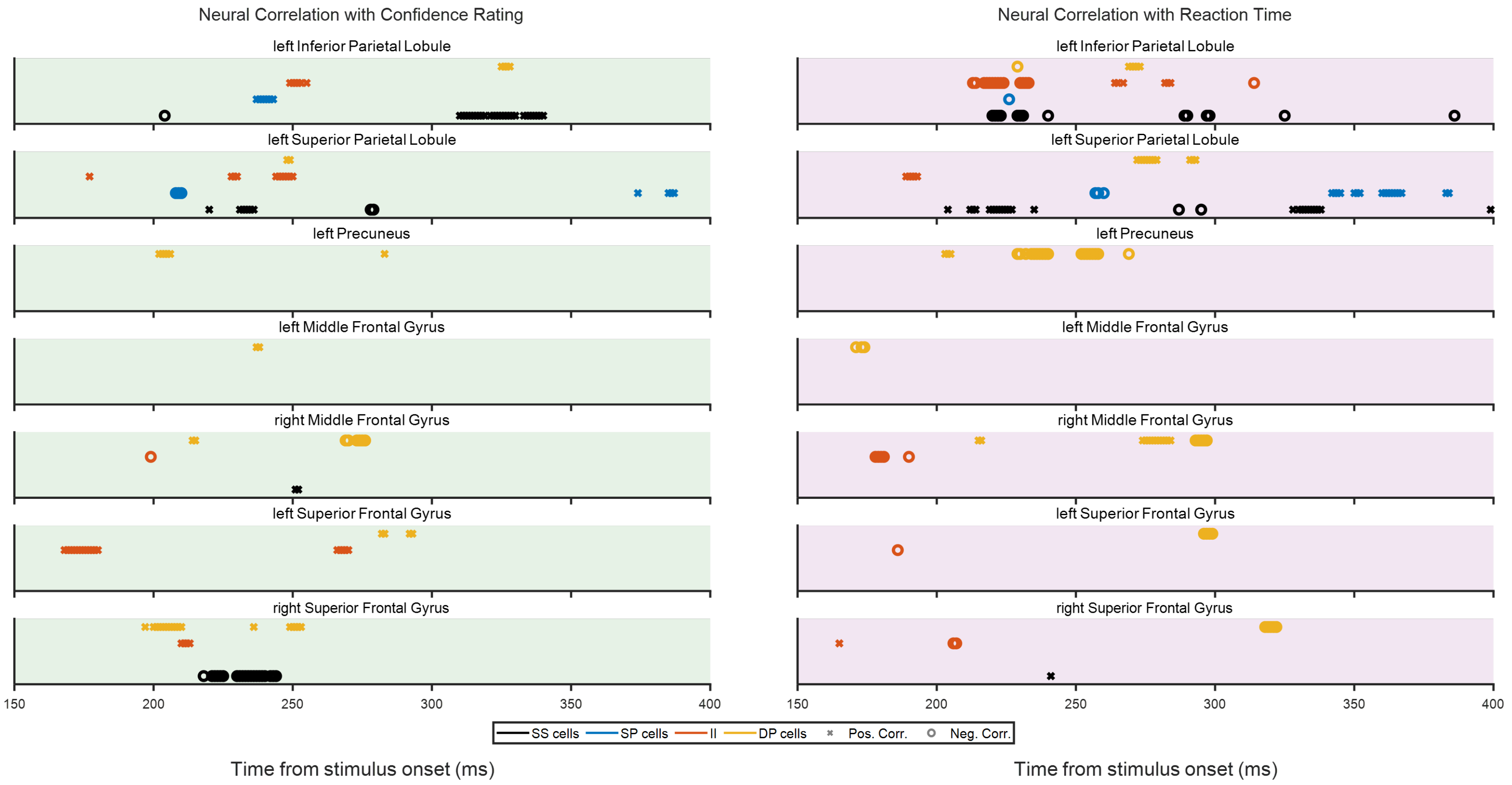


**Fig. S16. Correlations suggest SS activity in left IPL relate to subjective decision confidence and right SFG to decision uncertainty, with left PreCUN linked to faster choice-based RTs (objective confidence proxy) and left SPL to slower choice-based RTs (objective uncertainty proxy).** Correlation of estimated source activity (SS, SP, and DP and II neural populations) with subjective confidence ratings and choice-based RTs in high confidence trials, delineating the distinct roles of these regions in encoding cognitive aspects of confidence and decision-making speed. Trial-by-trial DCM analysis using support vector regression (SVR) reveals that SS activity in left IPL correlates with subjective confidence and those in the right SFG with uncertainty, while activities in the left PreCUN and the left SPL are associated with faster and slower RTs, respectively, hinting at their roles in objective confidence and uncertainty.

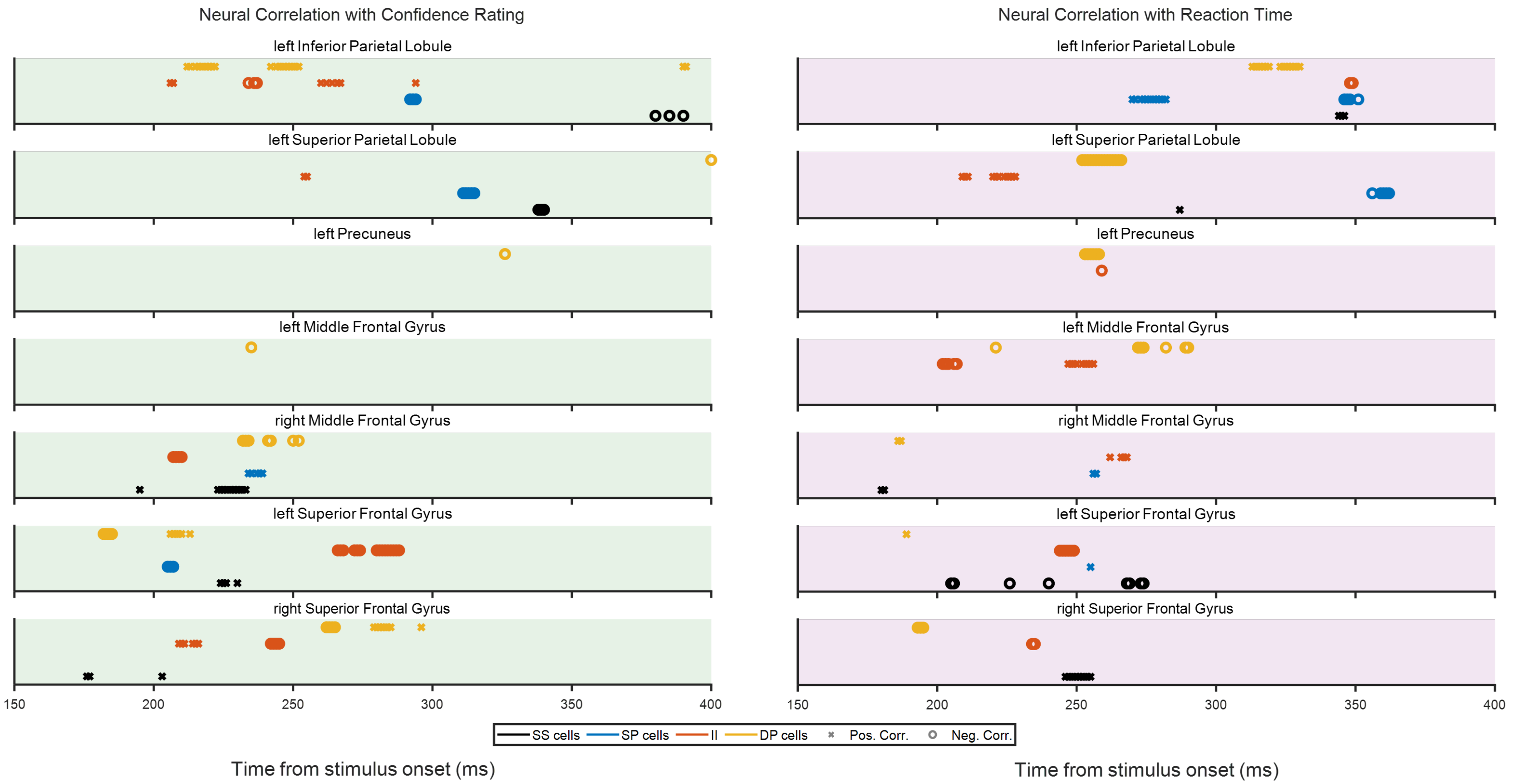


**Fig. S17. Estimated activities of II in the left SFG are associated with the encoding of subjective decision uncertainty in low confidence rating trials indicated by trial-by-trial DCM.** Correlation of estimated source activity (SS, SP, and DP and II neural populations) with subjective confidence ratings and choice-based RTs during low confidence rating trials, providing insights into the dynamic neural processes underpinning confidence and decision-making speed. Trial-by-trial DCM analysis using SVR reveals that II activity in left SFG correlates with subjective decision uncertainty.

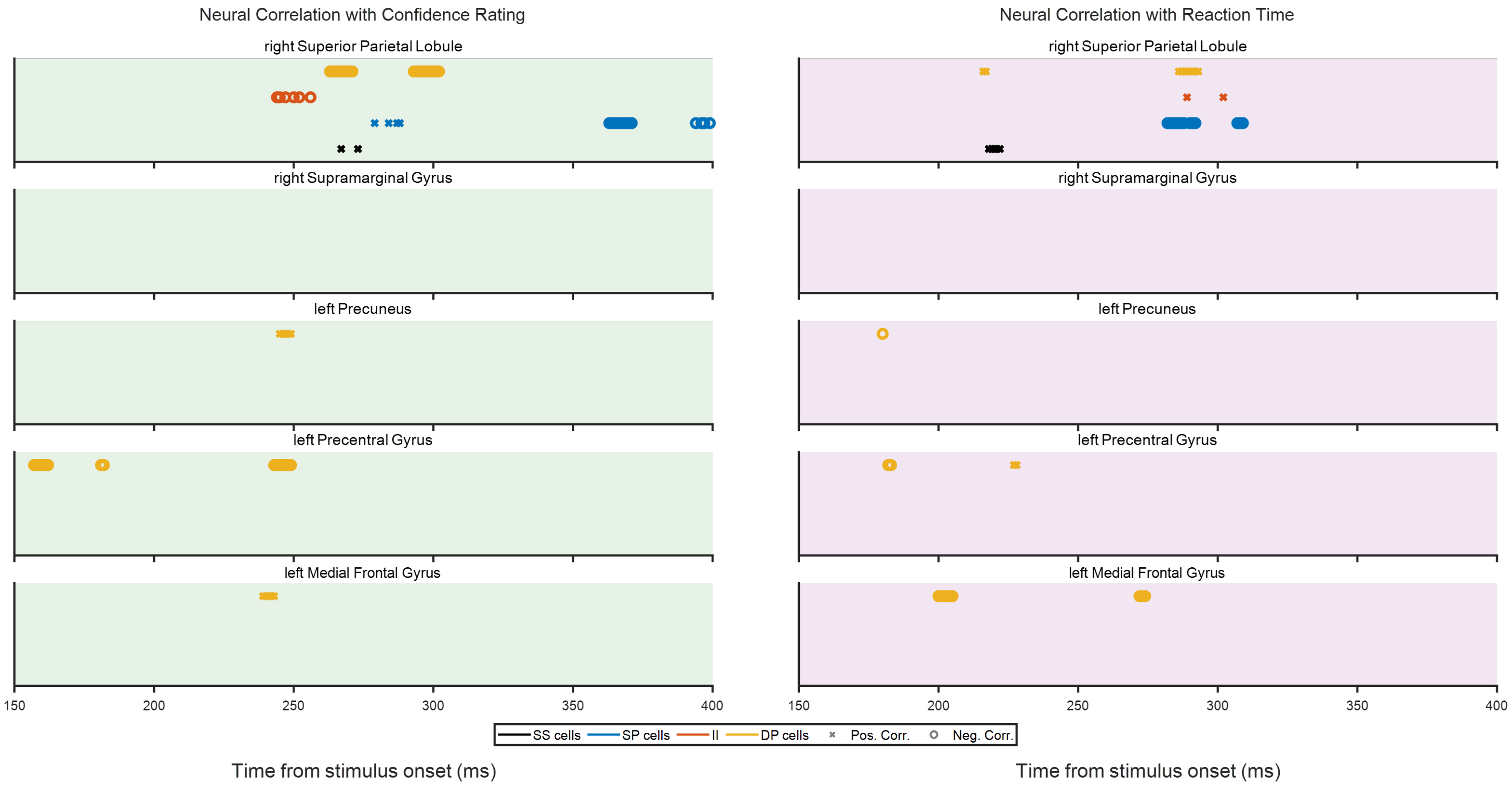


**Fig. S18. Estimated activities of SP and DP neural populations in right SPL correlate with subjective decision uncertainty.** Correlation of estimated source activity (SS, SP, and DP and II neural populations) with subjective confidence ratings and choice-based RTs, delineating the association neural activity within neural populations confidence levels and decision-making speed during trials with fast choice-based RTs. Trial-by-trial DCM analysis using SVR reveals that SP and DP neural population activity in right SPL correlates with subjective decision uncertainty.

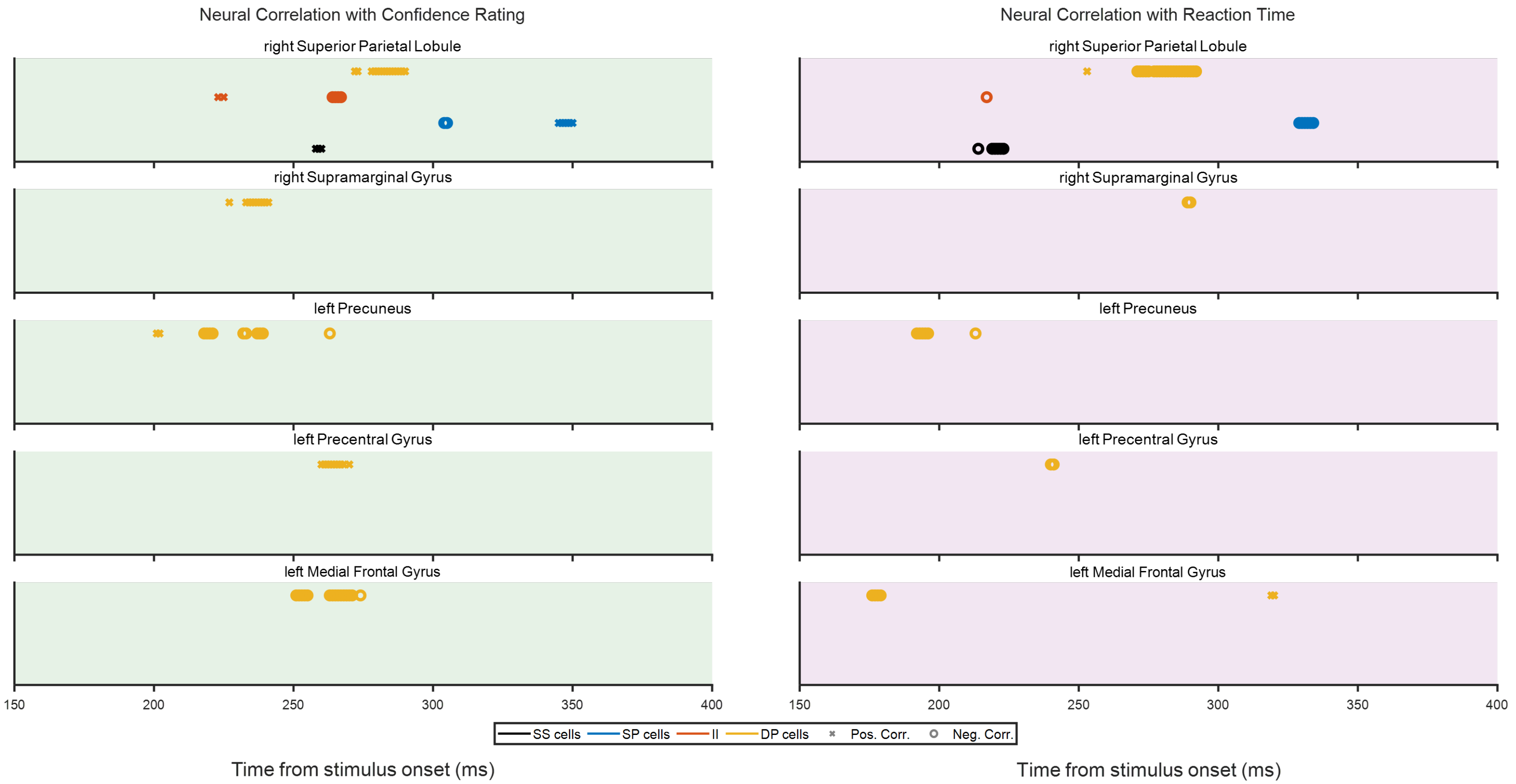


**Fig. S19. In slow RT trials, estimated DP activity in left PreCUN correlates with confidence rating, while estimated activities of DP neural poulation in right SPL correlates with faster choice-based RTs.** Correlation between estimated source activity and both subjective confidence ratings and choice-based RTs, during trials with slow choice-based RTs, highlighting the neural correlates of confidence and decision speed. Trial-by-trial DCM analysis using SVR shows that DP neural population activity in left PreCUN correlates with confidence rating suggesting its role in subjective decision confidence whereas DP activity in right SPL correlates with faster choice-based RTs hinting its role in objective decision confidence.
